# Enabling Skilled Human-Computer Interaction After Paralysis via a Wearable sEMG Interface

**DOI:** 10.64898/2026.01.09.698484

**Authors:** Dailyn Despradel Rumaldo, Max Murphy, Luigi Borda, Najja Marshall, Emanuele Formento, Mario Bräcklein, JinHyung Lee, Jun Ye, Peter Walkington, Dano Morrison, Stephanie Naufel, Kriti Kacker, Nikhil Verma, Jenn Shannahan, Mauricio Saavedra, Ahmed Gurafi, Pok Him Siu, Zaki Alam, Amy Boos, Jennifer L. Collinger, Diego Adrian Gutnisky, Douglas J. Weber

## Abstract

Most individuals with tetraplegia retain some myoelectric function in their forearms, which offers the possibility of using surface electromyographic (sEMG) control for human-computer interaction (HCI). We demonstrate the potential of this approach by showing that people with motor-complete *(n=5)* and motor-incomplete *(n=2)* tetraplegia can accurately control myoelectric activity in their forearm to perform discrete button-click and continuous positioning tasks. These control inputs were mapped to the firing rate of motor units detected by a wireless wristband sensor designed for everyday use. Participants completed four testing sessions to assess their speed and accuracy. Motor units that displayed a wide dynamic range in their firing rate performed best during tasks requiring continuous, single-axis control. Interestingly, the level of impairment did not affect performance on the clicking and 1D cursor control tasks. However, those with motor-incomplete injuries showed greater independent control over two motor units than participants with motor-complete injuries, who exhibited stronger coupling between units. Participants also confirmed the practical utility of the device, successfully placing and removing the sEMG wristband on their own and consistently rating it as comfortable and easy to manage. These findings are significant because they offer the first demonstration of motor unit-based control in individuals with cervical spinal cord injury (SCI) using a fully wearable wristband interface, highlighting the feasibility of moving these systems out of the lab and into daily life.

**One-Sentence Summary:** People with tetraplegia used a wristband sensor to detect forearm motor unit firing and perform human-computer interaction tasks.

## INTRODUCTION

Humans interact with various digital and robotic devices to accomplish essential activities of daily living. Although voice-based interactions are becoming increasingly useful for human-computer interaction (HCI) applications, manual interactions via keyboard, mouse, and other handheld controllers remain the dominant medium (*1*). However, people with tetraplegia lack the manual dexterity needed to use traditional input controllers, restricting their access to digital devices and limiting their ability to use assistive robotic arms (*2*). Thus, alternative HCI technologies are needed to restore autonomy and increase access to digital devices for people with tetraplegia.

Surface electromyography (sEMG) based wrist wearables have emerged recently as a viable HCI solution that can be performant at scale (*3*) and may even offer new promise for those with high-level (e.g., cervical) motor complete spinal cord injury (SCI). A growing number of studies demonstrate that a majority of people with motor complete SCI retain the ability to activate spinal motor neurons below the level of injury, offering the potential to use myoelectric controllers for computer-based tasks (*4–7*). Indeed, Ting et al. demonstrated that myoelectric and motor unit activity could be detected and classified to predict attempted movements of individual fingers in a person with motor complete tetraplegia (*6*). Souza-Oliveira demonstrated recently that people with a range of impairment levels after SCI could control an avatar to generate virtual hand gestures by volitionally modulating the firing rates of motor units in their forearm (*7*). While the potential for sEMG-based control for HCI is becoming apparent, it is unclear whether wearable devices can deliver the sensing capabilities required to enable crucial control functions for people with SCI.

In this study, we demonstrate that individuals with tetraplegia *(n=7)* due to cervical SCI can perform standard HCI tasks by controlling the firing rate of individual motor units, which were detected in real-time using a wireless neuromotor interface in a wristband form factor (*8*). Motor unit spiking activity was recorded as participants attempted to form prompted hand gestures, such as flexion or extension of specific joints. All participants were able to recruit and control multiple motor units for each of the prompted hand gestures, even those that were paralyzed. At the start of each session, a calibration was performed to generate a catalog of motor units associated with each hand gesture. A live display of the spiking activity was provided during calibration to verify that participants could control the motor unit firing rate before proceeding to the HCI tests. During HCI testing, participants were instructed to control the firing rate of 1 or 2 selected motor units by attempting the associated hand gesture. The firing rate for a selected motor unit was mapped to discrete (e.g., button-click) or continuous (e.g., cursor positioning) control inputs, and participants completed four testing sessions to assess their speed and accuracy in completing HCI tasks that required real-time modulation of the timing and rate of motor unit firing. The tests were designed to represent core primitives of modern HCI controls, specifically selection and navigation (*9*).

All participants were able to complete the HCI tests using motor units associated with a variety of hand gestures. Several factors influenced the speed and accuracy of their control, particularly during initial sessions. However, by the fourth session, all participants demonstrated proficient control of motor unit firing rates for generating discrete clicks and continuous cursor positioning, and their motor unit control was surprisingly unaffected by the impairment level for the associated gesture. Moreover, all participants successfully achieved independent control of two motor units for two-dimensional cursor movements. Participants also demonstrated that they could independently don and doff the sEMG wristband, consistently rating the device as useful, comfortable, and easy to manage. These findings highlight the potential of sEMG wristbands to enable rich, hands-free computer interactions for individuals with tetraplegia, supporting control of assistive devices and activities such as gaming, communication, and social engagement. Leveraging the technical capabilities of a system that is widely available and includes accessible controls for human-computer interactions would be an extremely powerful and practical option for enhancing accessibility.

## RESULTS

### Participants

The study was approved by the Institutional Review Boards at Carnegie Mellon University and the University of Pittsburgh. Nine individuals with tetraplegia from cervical SCI were enrolled, of whom two withdrew due to transportation and scheduling constraints. After providing informed consent, each participant underwent a physical examination to determine impairment level using the American Spinal Injury Association Impairment Scale (AIS), administered by a licensed Occupational Therapist. Five participants had motor-complete tetraplegia (AIS A: n = 4, AIS B: n=1) and two had motor-incomplete tetraplegia (AIS C: n=1, AIS D: n=1). Manual muscle testing (MMT) was performed to measure residual strength for the wrist and finger joints (Fig. S1). The cohort of 1 female and 6 male participants ranged in age from 21 to 72 years (median 37 years) and in time since injury from 2 to 22 years (median 10 years).

### Study Design

Seven participants completed one wristband screening session followed by at least four testing sessions, each lasting up to four hours. The screening session was performed to verify that sEMG and motor unit activity could be detected. Participants then completed four testing sessions in which they controlled the firing rates of individual motor units to generate discrete button presses or continuous cursor positioning tasks. These tasks assessed the participants’ ability to recruit, modulate, and derecruit spiking activity in motor units associated with different hand gestures. Performance was quantified primarily through task completion rate and accuracy. Although the study did not include an explicit training protocol, participants often demonstrated measurable improvements in control performance across sessions, reflecting learning and adaptation to the interface. Device usability was also assessed by observing participants’ ability to don and doff the wristband and by collecting usability feedback.

### Screening and calibration for motor unit detection

All tasks were performed with participants seated in their personal wheelchairs facing a display monitor (Fig. 1A). During the initial screening and calibration phase, the wristband was placed at multiple forearm locations between the ulnar head (at the wrist) and the lateral epicondyle (at the elbow) to identify the optimal site for detecting motor units. Wrist circumference was measured at each location to determine appropriate wristband sizing before placement (Table 1 and Fig. S2A). Participants attempted prompted gestures with the wristband at each site, and the location yielding the greatest number of gestures with go-prompt-modulated, high-quality motor unit templates was used for all testing sessions (Fig. S2).

**Fig. 1.**
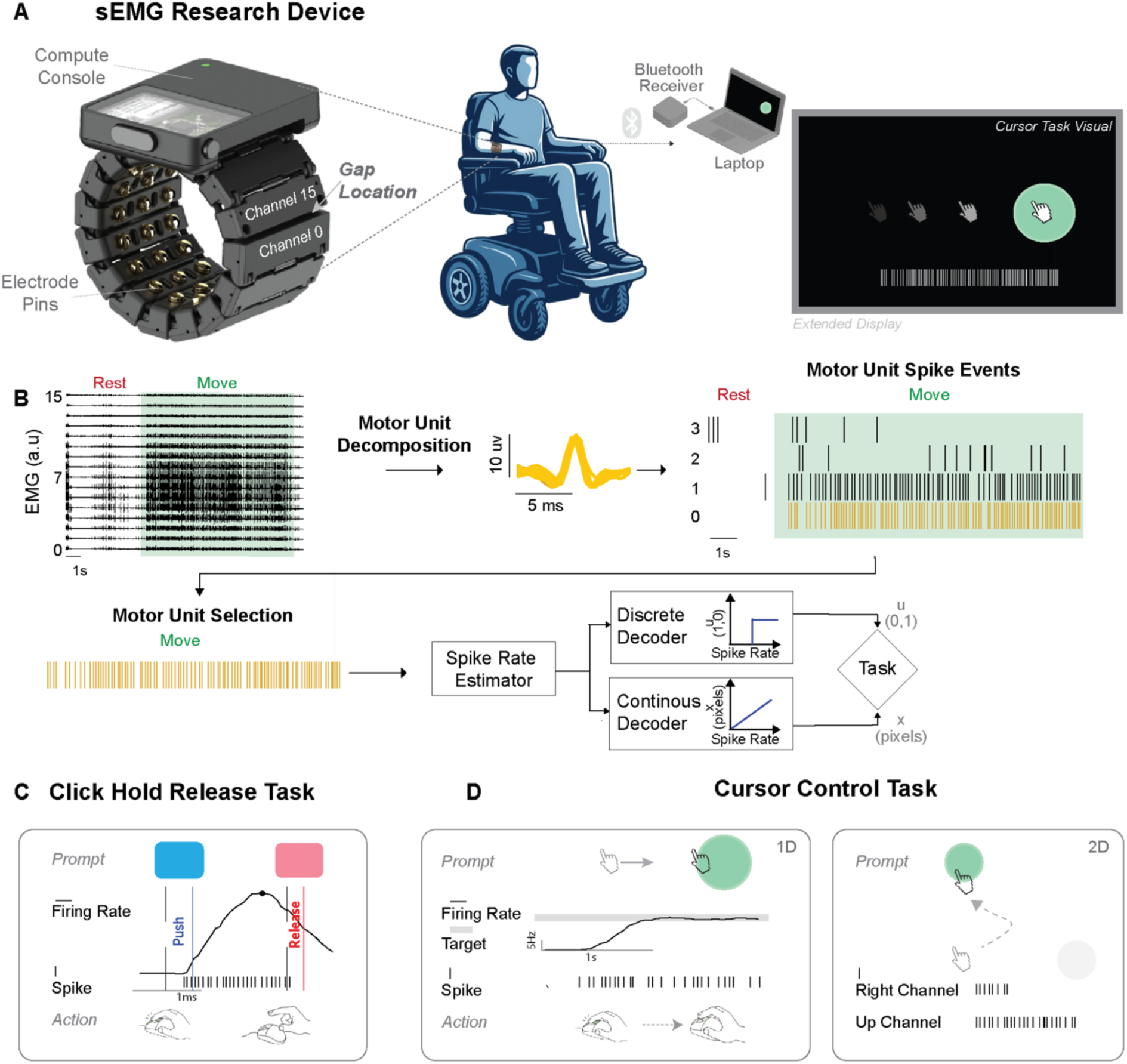
Experimental setup, calibration, and task framework. (**A**) The 16-channel sEMG wristband was positioned on one forearm of individuals with tetraplegia. The sEMG signals were transmitted via Bluetooth to a computer console and displayed on a screen for task feedback. (**B**) The sEMG signals were decomposed in real-time to identify motor unit action potentials (MUAPs). Individual motor units were selected for discrete and continuous task-control testing using the firing rate of the selected motor unit. (**C**) The Click-Hold-Release task required participants to increase and hold the motor unit firing rate above a threshold for a variable hold time and then release by reducing the firing rate. (**D**) Cursor control tasks were performed in one or two dimensions (1D, 2D). In the 1D task, participants modulated the firing rate of a single motor unit to move a cursor towards a target positioned along the horizontal axis. The 2D task required independent modulation of two motor units to reach targets along the horizontal (*Right Channel*) and vertical (*Up Channel*) axes.

**Table 1.** Participant information.

| Participant ID | Time Since Injury (years) | Sex | AIS | Injury Level | Band Size | Circumference at Device Site (cm) | Device Position* (% of forearm length) |
| --- | --- | --- | --- | --- | --- | --- | --- |
| MCP01 | 19 | M | A | C5 | L | 22 | 41.4 |
| MCP04 | 10 | M | A | C4-C5 | L | 20 | 48.3 |
| MCP05 | 22 | M | D | C4-C5 | XL | 23 | 68 |
| MCP06 | 14 | F | C | C6 | S | 14.5 | 54.2 |
| MCP07 | 5 | M | A | C4 | L | 20 | 38.5 |
| MCP08 | 2 | M | A | C5 | L | 18.5 | 48.1 |
| MCP09 | 7 | M | B | C6 | XL | 22 | 37 |

At the start of each session, participants completed a calibration procedure in which the models for detecting and classifying motor unit action potential (MUAP) were trained through real-time interactions using iterative fitting of the model parameters during alternating periods of rest and activity (Fig. 1B). Unlike the methods used for offline decomposition (*10*), this process was performed with the participant providing real-time volitional input and receiving visual feedback to facilitate stable detection of MUAPs (see example in Supplementary Movie 1). During this procedure, participants performed repetitions of an instructed gesture selected from a fixed list of possible gestures: pronation or supination of the forearm; flexion, extension, radial deviation, or ulnar deviation of the wrist; or flexion or extension of an individual digit. Participants were prompted to intentionally contract muscles related to the gesture. In a few cases, the prompted actions resulted in voluntary movement at the joint (e.g., wrist extension), and participants were instructed to limit their effort to avoid generating overt movement. A variety of metrics were used to select a specific motor unit for each gesture for the real-time control tasks. Briefly, these metrics quantified MUAP waveform isolation quality and controllability based on the alignment of motor unit spiking activity with movement epochs during alternating rest and movement sequences. We refer to the unique combination of gesture and associated motor unit as the gesture-motor unit pair.

The motor unit selected for each gesture (i.e., gesture-motor unit pair) was then used for real-time control in the discrete *Click-Hold-Release* (CHR; Fig. 1C) task and the continuous *1D* and *2D cursor control* tasks (Fig. 1D and Fig. S3). These complementary test modes reflect the standardized input categories recognized in the USB Human Interface Device (HID) specification (*11*). By aligning our task design with these established categories, we ensured that performance could be compared under paradigms that correspond directly to the principal classes of user input defined by international device standards.

### Motor unit control for discrete state-control tasks

In the *Click-Hold-Release* (CHR) task, participants were instructed to complete a fixed number of *Click*, *Hold*, and *Release* sequences using a virtual push-button (Fig. 2A). Participants controlled these interactions by activating or silencing the selected motor unit; *Click* actions were triggered by a single MUAP and release actions were triggered after MUAP firing lapsed for a minimum of 200 milliseconds (Fig. 1D, see example in Supplementary Movie 2). The required hold duration was varied between one and four seconds to prevent participants from anticipating the release timing.

**Fig. 2.**
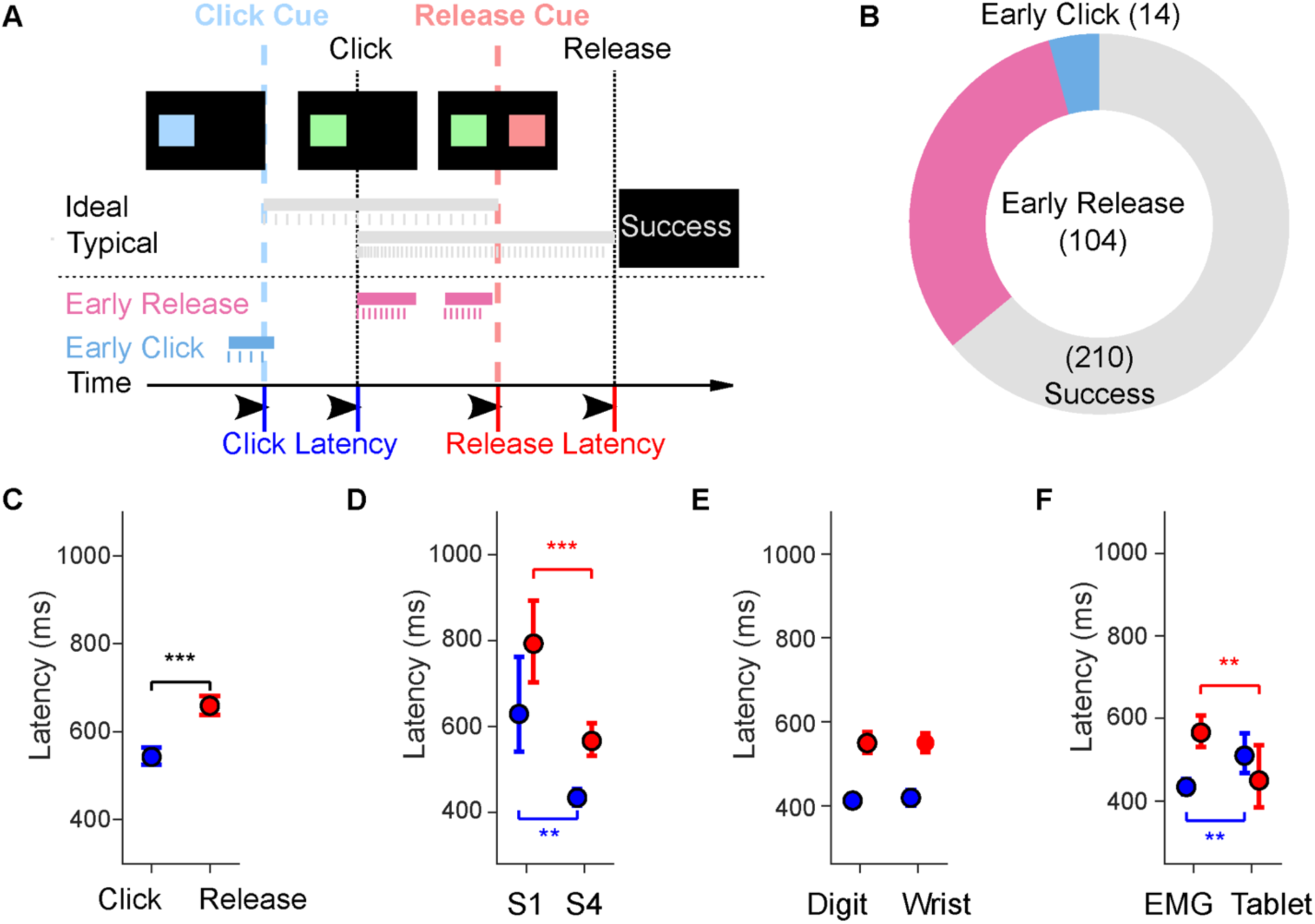
Push-button and Release (CHR) task design and performance across demographics and sessions. (**A**) A blue square (*Click* prompt) cued participants to generate a click event by recruiting and sustaining motor unit activity until the red square (*Release* prompt) appeared, prompting participants to deactivate the motor unit. Example timelines illustrate typical successful trials as well as early click and early release errors. (**B**) Distribution of best-case session outcomes across all participants in sessions requiring 30 completed trials, showing the proportion of successful trials (gray) versus early click (blue) and early release (pink) errors. (**C**) Click (blue) and release (red) latencies from all successful trials (across all session/gesture/subject combinations) using wristband control show a significantly faster average click latency (543 ms [526, 564 ms]), compared with the average release latency (659 [639, 682] ms). (**D**) Latencies grouped by AIS grade, plotted for Session 1 and Session 4 to assess training effects. Latencies decreased significantly from Session 1 to Session 4 for both actions. (**E**) Push and release latencies are further stratified by gesture group (wrist vs. digits). (**F**) Push and release latencies from S4 are shown for the wristband interface versus the tablet touchscreen. All error bars indicate the 95% confidence interval of the mean performance metric computed with bootstrap N=10,000. Statistical significance was assessed with two-tail bootstrapping (N=10,000): p<0.05 (*), p<0.01 (**), p<0.001(***).

As a primary task outcome, we measured the latency between the frame introducing the visual cue and its corresponding *Click* or *Release* event; a secondary outcome metric was task accuracy, which required participants to not only suppress activity at rest, but also to sustain firings of the selected motor unit with no longer than a 200 millisecond inter-spike interval. Across sessions, subjects completed the full click-hold-release sequence with a mean success rate of 52.9% (95% confidence interval [46.7%, 58.7%]). Subjects clicked responsively, with a mean sensitivity of 92.9% [89.9%, 95.6%], but failed to hold the firing rate above threshold on 26.9% of trials [22.9%, 31.0%]. Users consistently released the button when presented with the release cue (Fig. 2B).

On average, click events were generated with significantly lower latency than release events (543 vs 659 ms; 95-percentile difference of the means [-144, -87] ms, p < 0.001, using an N=10,000 two-tailed, centered two-sample bootstrap test of the mean difference (Fig. 2C). This difference is likely due to the requirement for the release, but not the click, that a minimum interval of 200-milliseconds must elapse to trigger the *release*. Latencies decreased significantly from Session 1 to Session 4 for both actions (click: 629 vs 434 ms, 95-percentile difference of the means 195 [103, 327] ms, p < 0.01; release: 792 vs 566 ms, 95-percentile difference of the means 227 [125, 335] ms, p < 0.001; Fig. 2D).

Surprisingly, latencies were not significantly different when grouping gestures as related to movements of the digits or wrist, despite the lower MMT scores associated with digits (Fig. 2E and MMT scores in Fig. S1).

We also tested the CHR task using a familiar tablet touchscreen as a point of comparison. Wristband-based click responses were significantly faster than the tablet (434 vs 510 ms; 95-percentile difference of the means [-133, -30] ms, p = 0.0062; Fig. 2F), while release responses were faster on the tablet (565 vs 450 ms, 95th-percentile difference of the means 119 [24, 195] ms, p = 0.0082; Fig. 2F).

### Motor unit control for 1D cursor positioning

To evaluate continuous positioning control, participants first performed a task in which they controlled the firing rate of a selected motor unit to move a cursor to and hold at a target. The initial (resting) position was defined as the center of the screen, and the movement direction was assigned based on participant handedness and the natural movement direction of the selected gesture (e.g., wrist extension for a right-hand participant moved the cursor to the right on the screen). Task difficulty was varied by adjusting the target diameter and distance from the origin (Fig. 3A; see Methods).

**Fig. 3.**
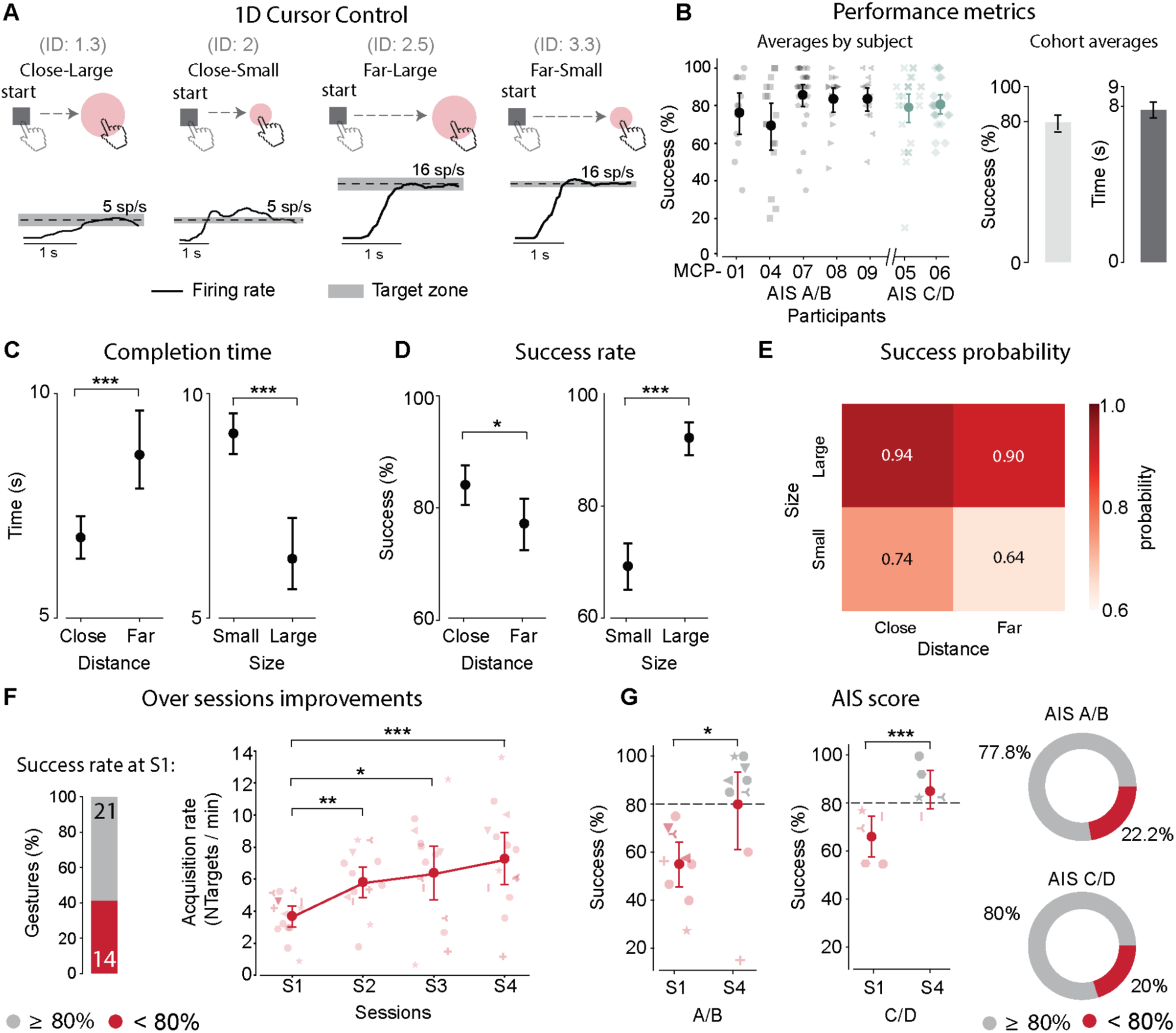
Performance in a one-dimensional control task using motor unit-based firing rate modulation. (**A**) Examples of the 1D control task, which included four target conditions defined by target distance (Close vs. Far) and size (Small vs. Large). The numbers in parentheses indicate the corresponding index of difficulty (ID) for each condition. Participants modulated the firing rate of selected motor units to reach and hold in the target zone, with targets requiring either low or high firing rates. Representative firing rate trajectories are shown in black for MCP08 while reaching the 4 targets; the target firing rate and zone are indicated by a dashed line and shaded area, respectively. (**B**) Quantification of the success rate across participants and average performance (success rate and reach completion time) for all the participants and gesture-motor unit pairs (N = 7 participants x 4 sessions). Quantification of the effects of target size and distance on (**C**) success rate and (**D**) completion time. (**E**) Heatmap reporting success probability based on target size and distance. (**F**) *Left*: bar plot reporting the percentage of gesture-motor unit pairs with a success rate below (gray) and above (red) 80% at session 1. *Right*: quantification of changes in acquisition rate across sessions for the subthreshold gesture-motor unit pairs (< 80% success rate). (**G**) *Left*: comparison of the success rate at session 1 (S1) and session 4 (S4) for motor complete (AIS A/B) and incomplete (C/D) participants. Right: percentage of gesture-motor unit pairs with success rate above (gray) and below (red) 80% at S4 for motor complete and incomplete participants. The circle marker on the plots indicates the mean value of the performance metric. All error bars indicate the 95% confidence interval of the mean performance metric computed with bootstrap N=10,000. Statistical significance was assessed with two-tail bootstrapping (N=10,000): p<0.05 (*), p<0.01 (**), p<0.001(***).

Cursor position was mapped linearly to the smoothed firing rate of the selected motor unit. The firing rate was normalized to each participant’s maximal steady-state firing rate (SSFR), defined as the highest firing rate participants could sustain comfortably during a brief preparation period. To complete a trial, participants were required to increase the firing rate of the selected motor unit to position the cursor over the target zone for 1s, holding the firing rate within a small or large range, indicated by the target size. A trial was considered successful if the cursor acquired the target before a fixed 15s timeout. Supplementary Movie 3 explains the task and contains examples of successful and failed trials.

Each participant completed at least four sessions, and Fig. 3B shows the average success rate for each participant across all gesture-motor unit pairs and sessions. Overall, participants achieved an average success rate of 79.6%, with a mean completion time of 7.79 s. To assess the impact of task difficulty, we quantified success rate and completion time across the four target conditions for all the sessions and gestures tested (see summary table in Fig. S3A). Performance declined with increasing target distance and decreasing target size (Fig. 3C, D). Specifically, completion times were longer for targets that were far compared to close targets (8.6s vs 6.8s, 95th-percentile difference of the means [0.92, 2.92], p<0.001), and for small targets compared with large targets (9.1s vs 6.3s, 95th-percentile difference of the means [1.75, 3.63], p<0.001). Success rates were lower for far versus close targets (77.2% vs 84.1%, 95th-percentile difference of the means [-12.8,-1.11], p<0.05) and for small versus large targets (69.2% vs 92.1%, 95th-percentile difference of the means [-27.9,-17.9], p<0.001). Interestingly, despite the nominally matched index of difficulty (ID), computed using the Fitts’ Law formulation (*12*) between the far-large target (2.46 bits) and the small-close target (2.0 bits, Fig. 3A), participants were significantly more successful completing trials for the far-large target, despite the greater firing rate required (89% vs 77%, 95th-percentile difference of the means [6.7, 18.3]%, p<0.001; Fig. 3E).

We also quantified changes in the target acquisition rate from sessions 1 to 4. Because many participants achieved ≥80% success rates with some gesture-motor unit pairs in session 1 (Fig. S3A), the analysis was limited to gestures yielding success rates below this threshold to examine their potential for improvement (Fig. 3F; n = 14 gestures with success rate < 80%, compared to n = 21 with success rate ≥ 80%). Within this subset, acquisition rate improved significantly from 3.7 targets/min in session 1 to 7.3 targets/min in session 4 (Fig. 3F; 95th-percentile difference of the means [1.88, 5.3], p<0.001). Finally, we investigated whether impairment severity influenced performance gains. We found that both motor complete (AIS A/B) and motor incomplete (AIS C/D) participants improved their success rates from session 1 to 4 (Fig. 3G). The proportion of participant–gesture pairs exceeding 80% success increased comparably across groups: 77.8% and 80.0% of initially subthreshold gestures (i.e., success rate < 80%) exceeded 80% at session 4 for the motor complete and incomplete group, respectively (Fig. 3G). Although performance scaled inversely with task difficulty, participants showed significant improvements with training, regardless of impairment severity.

### Motor unit control for 2D cursor positioning

After establishing reliable 1D cursor control, we next examined whether participants could extend single motor unit control to 2D cursor movements. Gesture-motor unit pairs that achieved a success rate of ≥80% in the 1D control task (Fig. S2A) were considered eligible for testing in the 2D cursor control paradigm. When more than two gestures exceeded this 80% threshold, we evaluated as many pairwise combinations as permissible within the session duration without causing fatigue or discomfort. In this task, the firing rates of two selected motor units were mapped separately onto the horizontal and vertical coordinates of the cursor, enabling control along two orthogonal axes (see examples in Supplementary Movie 4). A single target appeared along either the horizontal or vertical axis, with four difficulty levels defined by target distance and size (Fig. 4A).

**Fig. 4.**
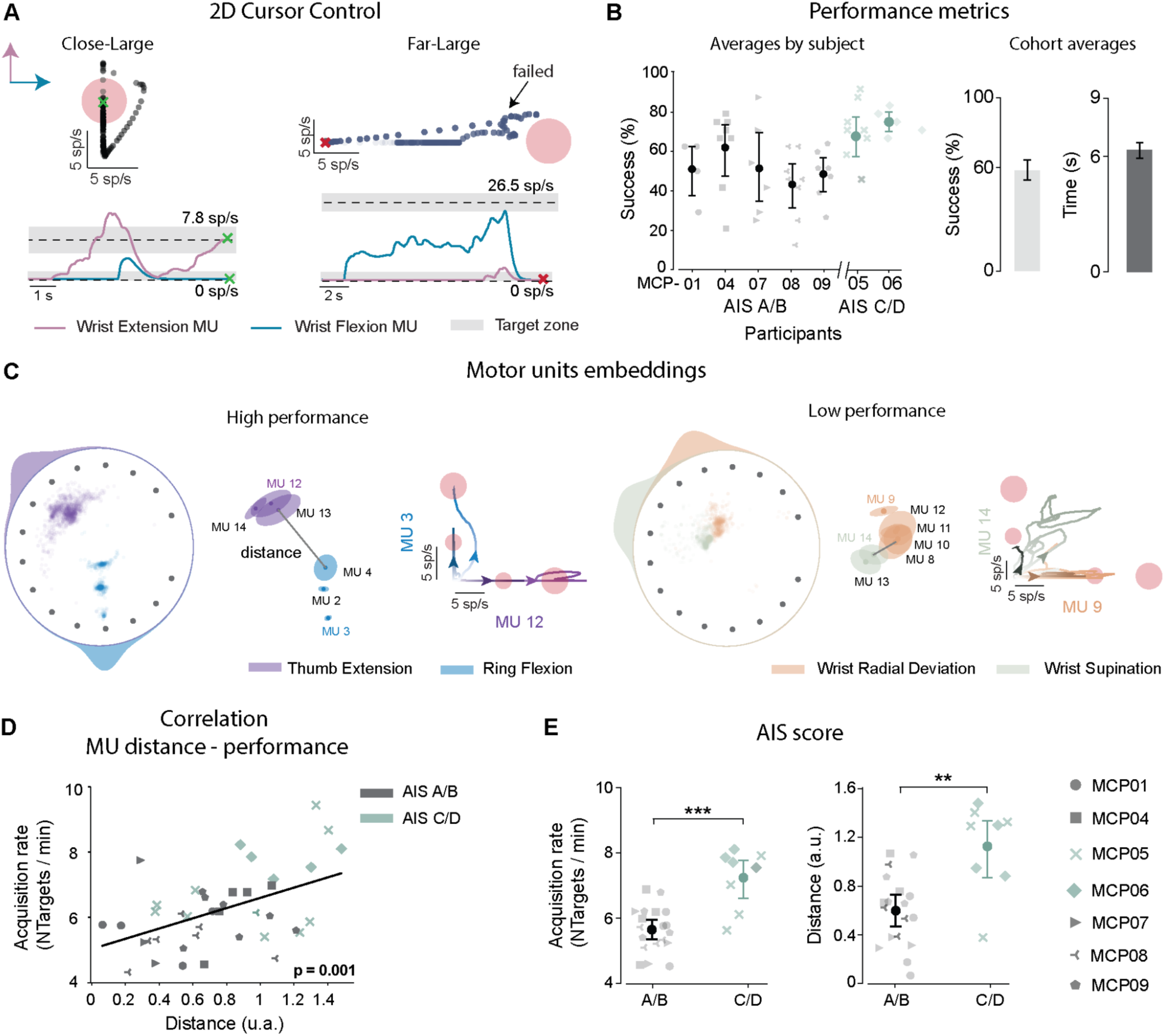
Performance on a two-dimensional cursor control task using motor unit firing rates for rate-to-position control. (**A**) Example of firing rate dynamics for 2D cursor control. The participant (MCP09) controlled one motor unit associated with wrist extension (pink) and another associated with wrist flexion (blue) to move the cursor along the vertical and horizontal axes, respectively. Green and red crosses indicate final firing rate values for a successful (left) and a failed (right) trial, respectively. (**B**) *Left*: success rate for each participant. *Right*: average success rate and completion time for all participants and sessions (N = 7 participants x 4 sessions). (**C**) *Left*: example of high task performance (> 80%), where motor units associated with distinct gestures (e.g. thumb extension, ring flexion) occupied well-separated regions of the embedding space, and firing rate trajectories of unit pairs showed clear task-related modulation. *Right*: example of low task performance (<80%), where motor units were less separated in the embedding, and their firing patterns overlapped across tasks. The green and red crosses indicate successful and failed trials, respectively. (**D**) Linear correlation between minimum L1 distance among recruited unit pairs with target acquisition rate (black line, linear fit; p = 0.001). (**E**) *Left*: Participants with AIS A/B (motor complete) achieved lower acquisition rates compared to participants with AIS C/D (motor incomplete). *Right:* quantification of the mean minimum L1 distance among motor unit pairs for motor complete and motor incomplete participants. The circle marker on the plots indicates the mean value of the performance metric. Error bars indicate the 95% interpercentile range of the mean performance metric computed with bootstrap N=10,000. Statistical significance was assessed with two-tail bootstrapping (N=10,000): p<0.05 (*), p<0.01 (**), p<0.001(***).

As expected, the 2D task was more difficult than the 1D task as it required participants to activate one motor unit without activating the other. Across all subjects and trials, the average success rate was 58% with a mean completion time of 6.3 s (Fig. 4B). To assess the effects of target distance and size, we analyzed two performance metrics: success rate and cumulative independent firing time (CIFT), which quantifies how effectively participants achieved independent control of the two motor units (*13*). CIFT values range from 0 to 1, where 1 indicates maximal independence. Consistent with the 1D results, the success rate varied with target distance and size (Fig. S3B, C). Specifically, success rates were lower for far versus close targets (48.3% vs 70.1%, 95th-percentile difference of the means [-31.4, -12.3], p<0.001) and for small versus large targets (50.8% vs 70.0%, 95th-percentile difference of the means [-25.5, -8.7], p<0.001) (Fig. S3B). However, only the target distance significantly affected the ability to independently modulate the two motor units. Far targets were associated with a significantly lower CIFT compared with close targets (0.59 vs 0.72, 95th-percentile difference of the means [0.08, 0.18], p < 0.001) (Fig. S3B), whereas no difference was observed between small and large targets (0.66 vs 0.65, 95th-percentile difference of the means [-0.04, 0.06], p = 0.62) (Fig. S3C).

We next examined whether the anatomical proximity of the selected motor units influenced task performance. For each gesture pair, we quantified the L1 distance between the unit-disk embeddings of the identified motor units (Fig. 4C), as computed when rendering visual feedback to the participants during motor unit model-fitting. We compared this distance with the average target acquisition rate for each session. We found a significant positive correlation between target acquisition rate and the minimum L1 distance (R = 0.49, *p* = 0.001; Fig. 4D), indicating that motor units located closer together were more difficult to control independently, likely due to greater shared common input (*14–16*).

We then assessed whether impairment severity affected 2D control performance. In contrast to the 1D task, motor complete (AIS A/B) participants exhibited lower performance compared with motor incomplete (AIS C/D) participants (5.6 vs 7.2 targets/min, 95th-percentile difference of the means [-2.22, -0.89], p<0.001). We also found that the minimum L1 distance between selected motor units was smaller in the motor complete group than in the motor incomplete group (0.6 vs 1.1, 95th-percentile difference of the means [-0.78, -0.23], p<0.01). Despite the lower performance compared to the 1D task, these results indicate that participants can control two motor units independently and simultaneously, particularly when the motor units are associated with anatomically distinct muscles. Thus, 2D positioning control is feasible using multiple motor units, even in people with severe tetraplegia.

### Motor unit firing rate properties during online control tasks

To gain insight into the neurophysiological factors that might underlie the performance differences observed across tasks and participants, we analyzed the relationship between motor unit firing rate characteristics and task performance. This analysis revealed that the ability to control a given motor unit depended critically on the dynamic range of the motor unit’s firing rate, which is limited by the maximal SSFR that the motor unit can sustain.

In the 1D cursor control task, success probability increased monotonically with SSFR across all participants (Fig. 5A). It was expected that success rates would be lower for the far targets, because they required higher firing rates to reach (Fig. 3D, E). However, this effect may be related to intrinsic properties of motor units, such as the motor unit type, that influence their tendency to exhibit lower or higher firing rates (*17*). To explore this further, we modeled individual trial success likelihood as a function of both the SSFR associated with the controlling motor unit, and the distance to the target (Fig. 5B). A logistic regression model including both normalized target distance and SSFR revealed that higher SSFR significantly improved success probability (β₂ = 0.076 ± 0.027, *p* = 0.005), whereas the negative effect of normalized target distance was weaker and only marginally significant (β₁ = −1.19 ± 0.63, *p* = 0.057). The interaction between normalized target distance and SSFR was not significant (β₃ = 0.034 ± 0.038, *p* = 0.37), indicating that SSFR exerted a broadly consistent stabilizing effect across both near and far targets. Thus, the likelihood of success was lower for far targets, but generally higher for units with a higher SSFR.

**Fig. 5.**
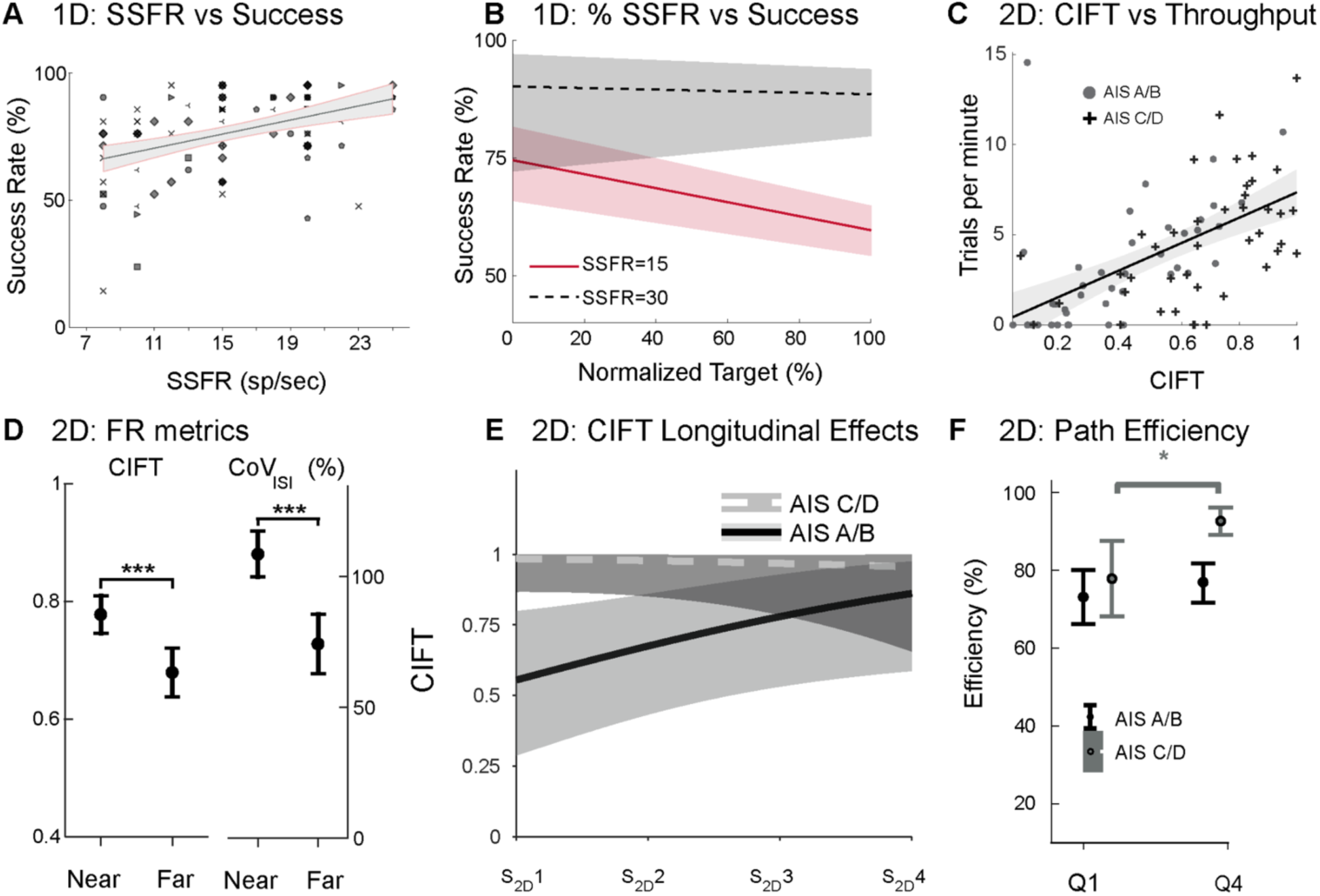
Motor unit firing-rate properties influence task performance. (**A**) Higher SSFRs are associated with higher success rates across trials in the 1D cursor task (R = 0.43, *p* < 0.001). The shaded area represents the t-based 95% confidence interval for the fitted line. (**B**) Predicted success rates in the 1D cursor task as a function of normalized target level for units with low (15 spikes/s) vs. high (30 spikes/s) SSFR, demonstrating improved control at higher tonic rates. (**C**) In the 2D cursor task, across participants, cumulative integrated firing time (CIFT) positively predicts trial throughput (trials per minute). (**D**) CIFT and CoV_ISI_ are significantly higher for near (easier) versus far (harder) targets. (**E**) Session-wise mixed-model estimates of CIFT indicate a significant and consistent improvement across training sessions, specific to the motor complete (AIS A/B) group. (**F**) Mean Path efficiency comparison between the first 25% (Q1) and final 25% (Q4) of trials within each set of 2D task trials, grouped by motor complete (AIS A/B, n=5) or motor incomplete (AIS C/D, n=2). Only the motor incomplete group demonstrated significant within-session improvements (79% vs 93%, t = 22.35, p = 0.028) in path efficiency. Except for panels B, C, and E, which use linear mixed effects, error bars indicate the 95% confidence interval of the mean performance metric computed with bootstrap N=10,000. Statistical significance was assessed with two-tail bootstrapping (N=10,000): p<0.05 (*), p<0.01 (**), p<0.001(***).

To quantify the relationship between MU independence and behavioral performance in the 2D cursor task (Fig. 5C), we fit a linear mixed-effects model predicting trial throughput (trials per minute) from the mean cumulative integrated firing time (CIFT) for each gesture–session combination, with a random intercept grouped by subject. The model revealed a robust positive association between CIFT and 2D task throughput (β_5_ = 6.29 ± 0.91, p<<0.001); a 0.15-unit increase in mean CIFT (on a 0–1 scale) corresponds roughly to an expected increase of approximately 1 trial/min. The subject-level random intercept variance was estimated at effectively zero for all participants, indicating that differences in baseline throughput across participants were negligible after accounting for CIFT. This result supports the hypothesis that good performance on the 2D task is produced, at least in part, by the ability to independently activate both motor units in the selected pair. As expected, CIFT decreased significantly when participants attempted to reach targets that were further away (0.72 [0.69, 0.76] for near vs 0.59 [0.56, 0.63] for far targets, p < 0.001; Fig. 5D, *left*). There was also a significant decrease in the coefficient of variation for the interspike intervals (CoV_ISI_) while holding in the final second of successful trials, for the far targets compared to the near ones (54.8% [49.0%, 60.7%] for near vs 47.8% [43.3%, 52.1%] for far targets, p < 0.001; Fig. 5D, *right*).

In total, there were 46 different 2D gesture-session combinations (median: 7 gesture-session combinations/subject; min: 2 gesture-session combinations, MCP01; max: 10 gesture-session combinations, MCP09). Because of the inter-subject differences in gesture-motor unit associations, total number of available motor units, and performance in the 1D task by session and subject, it was not possible to model learning or practice effects for each gesture pair. Encouragingly, in our mixed-effects analysis, we saw an overall trend when grouping across all pairs of gestures such that participants tended to increase their ability to independently engage motor units with repeated practice across sessions (Fig. 5E). CIFT increased significantly across sessions (β_8_ = 0.47 ± 0.23, *p* = 0.042), indicating that participants tended to improve their independent control across repeated 2D-cursor practice sessions. As expected, participants with motor-incomplete injuries (AIS C/D) showed substantially higher baseline CIFT than those with motor-complete injuries (AIS A/B: β_9_ = 1.32 ± 0.60, *p* = 0.028). The interaction between the model’s coefficient for session and categorical grouping for motor complete/incomplete trended negative (β_10_ = -0.71 ± 0.40, *p* = 0.077), suggesting that AIS C/D participants, who already demonstrated relatively high CIFT in the earlier sessions, exhibited smaller gains across sessions, whereas several AIS A/B participants exhibited marked improvements over time (see Table S1). Analysis of the path efficiency measurements revealed that participants in the motor incomplete group (AIS C/D) demonstrated a significant within-session increase in efficiency from the first to the last quartile of trials (mean_Q1 = 79.01%, mean_Q4 = 92.97%; Δ = 13.97%; p = 0.028), indicating straighter trajectories as the session progressed (Fig. 5F). In contrast, participants with motor complete injuries (AIS A/B) showed no significant change across quartiles (mean_Q1 = 71.82%, mean_Q4 = 74.70%; Δ = 2.88%; p = 0.687). Together, these results highlight that motor-incomplete individuals exhibit rapid within-session refinement of control strategies, whereas motor-complete individuals show more gradual but larger improvement across sessions.

### User impressions and usability

This sEMG device is unique in that it enables real-time detection of motor unit firing in a lightweight and untethered form factor that can be donned quickly without requiring wet electrodes or skin preparation. Given its wearable form factor and potential for daily use, we gathered participant feedback via survey questions about the comfort, convenience, performance, and practicality to evaluate the potential for integration into daily routines. Across all four categories, responses are predominantly clustered at ratings 4 and 5, indicating the participants consistently found the system comfortable, straightforward to set up, reliable in performance, and practical for routine use. Median ratings on the 5-point Likert scale were 4.0 for Comfort, Convenience, Performance, and Practicality (Fig. 6A).

**Fig 6.**
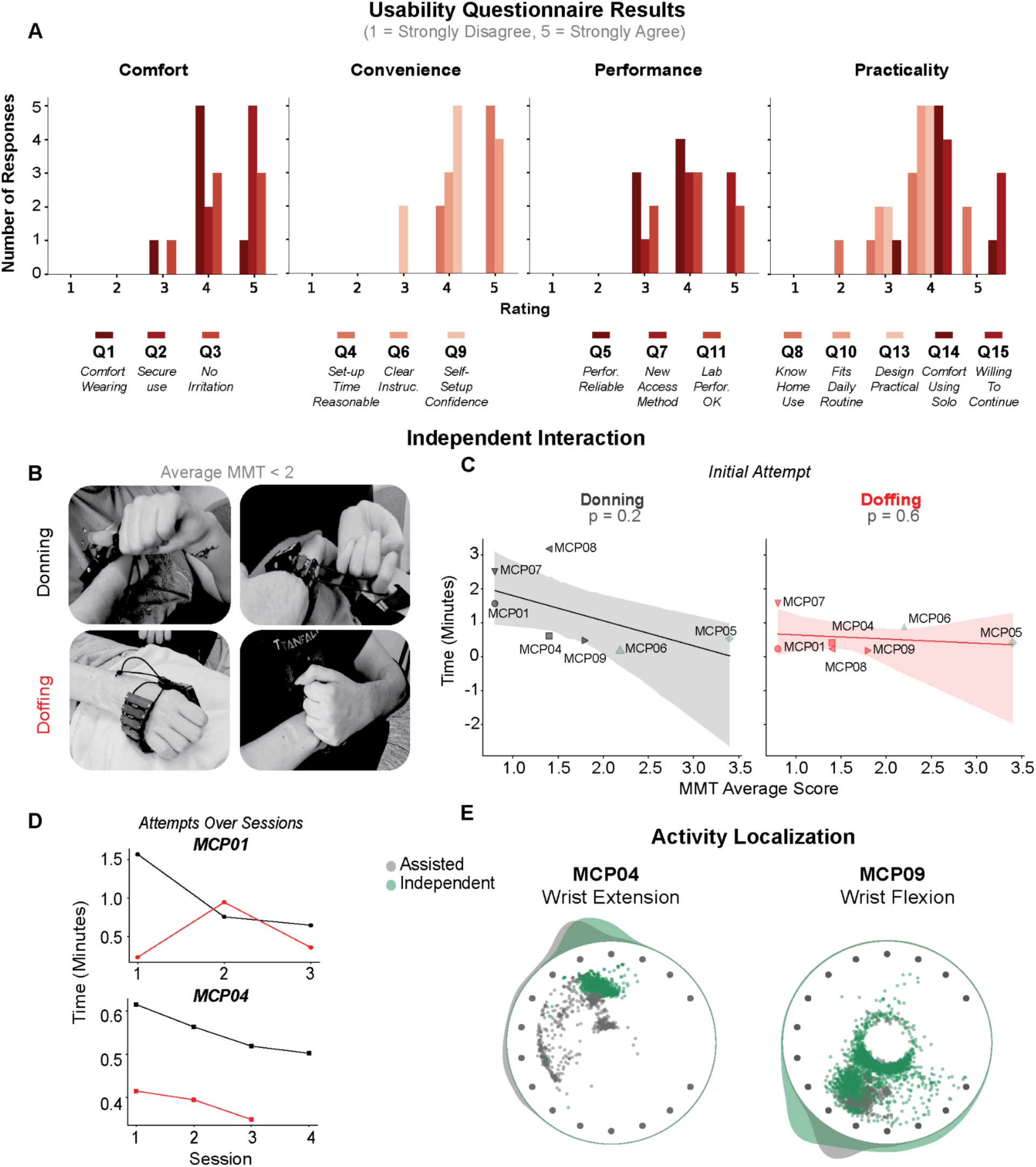
User feedback and usability assessment of the wearable sEMG system. (**A**) Participants completed surveys to provide user feedback in four categories: *Comfort*, *Convenience*, *Performance*, and *Practicality*. The histograms show the distribution of participants’ ratings (x-axis: rating from 1-5; y-axis: number of responses) for each question, grouped into the four categories. (**B**) Example donning and doffing strategies used by AIS-A participants. (**C**) Donning *(left)* and doffing *(right)* times as a function of average manual muscle test *(MMT*) score. Shaded regions represent 95% confidence intervals. (**D**) Donning time across multiple sessions for two representative participants (MCP01 and MCP04), showing reduced setup time with repeated use. (**E**) Electrode layouts (black dots) show motor unit activity during Independent (red) and Assisted (gray) trials for MCP04 (Wrist Extension) and MCP09 (Wrist Flexion). The motor unit embeddings for the assisted and unassisted trials show overlapping clusters, indicating that motor unit activity was co-localized across the setup conditions.

We also conducted a series of donning and doffing sessions, ensuring each participant attempted the donning and doffing sequence at least once (Fig. 6C). Results indicate that donning time was weakly correlated with residual upper-limb muscle strength as measured by MMT scores *(r = 0.58),* with less impaired participants donning the device more quickly. The correlation between doffing time and impairment level was even lower (*r* = 0.22), likely due to the simpler nature of the task. Participants employed creative strategies, such as using nearby surfaces to remove the device or positioning the band using their fingers, to complete the setup efficiently (Fig. 6B, see Supplementary Movie 5. Calibration was performed separately for the assisted and independently donned sessions, yet motor units could be detected reliably across repeated donning and removal sequences, with activity localized to similar channel regions in both conditions (Fig. 6E). While the wristband was not designed for use by people with impaired hand function, our findings demonstrate strong potential for independent use. With further design refinements, such as self-adjusting straps, the system could become a viable tool for out-of-lab applications and broader accessibility in everyday environments.

## DISCUSSION

Human-computer interaction (HCI) has become central to everyday life with the integration of digital systems in virtually all aspects of modern society, including the workplace, healthcare, education, communication, and entertainment. Manual HCI technologies remain critical in modern society, because they provide precise, reliable, and low-latency interaction between humans and digital systems. However, standard HCI technologies are difficult or impossible to use for the nearly 200,000 people in the United States living with tetraplegia due to SCI (*18*), posing additional restrictions on their access to important aspects of modern society. This unmet need has motivated the development of neuromotor interfaces that permit users to generate HCI control inputs through modulation of neural activity, as detected by sensors placed near the brain (*19*, *20*) or on muscles (*5*, *6*, *21*), thus enabling expression without physical action. While the potential benefits of these technologies have been demonstrated, access and adoption remain low.

Here, we demonstrate that a wearable wristband sensor can support crucial HCI control functions in people with tetraplegia due to SCI. The wristband sensor is untethered and can be donned and doffed quickly, without requiring skin preparation of conductive gel. We found that the sensor enables reliable detection of motor unit action potential firing in forearm muscles, and that users could modulate the firing rate of motor units associated with several different attempted movements of the wrist and fingers. Five of the seven participants in this study had motor-complete SCI (AIS A/B), yet they could proactively recruit and modulate the firing rate of motor units in muscles that were below the level of injury. These results are consistent with other studies (*5*, *6*, *21*) demonstrating the persistence of myoelectric function in people with severe paralysis, which is sometimes referred to as discomplete SCI and indicates partial sparing of neuromotor circuits (*22*).

The results presented here demonstrate that HCI control of discrete and continuous actions can be achieved with a compact, comfortable, and user-friendly wearable interface, thereby bridging the gap between laboratory systems and practical, everyday myoelectric control. The speed and accuracy of control achieved by our participants is on par with results reported in similar studies using research-grade EMG technologies that use hydrogel electrodes (*21*, *23*). Such systems are not designed for daily use and cannot be donned without assistance.

### Factors influencing controllability

All participants were able to control motor unit activity to perform discrete and continuous control tasks, but their speed and accuracy varied widely across the pool of motor units that were tested. The steady-state firing rate (SSFR) of individual units emerged as a consistent predictor of motor unit controllability. Units with higher SSFR enabled better performance in both discrete (e.g., push-button) and continuous (1D and 2D positioning) tasks. This finding suggests that the presence of intact, high-SSFR motor units could serve as a physiological benchmark for identifying viable control channels before real-time testing. Indeed, the recruitment threshold, firing rate modulation, and other intrinsic properties of motor neurons influence their behavior during voluntary control of muscle force (*24*) and should be considered when selecting motor units for HCI control tasks. Furthermore, motor neuron properties adapt to training (*25*) and may improve over time.

Additional factors are important to consider for HCI tasks that require controlling multiple degrees of freedom (DoF). The 2D cursor positioning task used here required participants to simultaneously and independently control the firing rate of two motor units. All participants were able to perform the 2D control task, but their performance was better when the selected motor units were more spatially separated (Fig. 4C). We did not consider anatomical proximity when selecting motor unit pairs, but functional independence of motor units is an important factor to consider when selecting motor units for multi-DoF control. Furthermore, training exercises may be effective in improving controllability, even for motor units in the same muscle (*13*).

### Effects of Impairment Severity

Across the participant pool, injury levels ranged from C4 to C6 and spanned AIS grades A through D. Despite this range, all were able to voluntarily modulate spinal motoneurons controlling wrist and finger movements. Notably, the motor pools innervating these muscles are located primarily within the C5–C7 segments (*26*, *27*), and thus likely below the lesion level for most participants in our cohort (n = 5 with injuries at C4–C5 and n = 2 at C6, see Table 1). Nevertheless, all participants achieved accurate 1D and 2D cursor control, demonstrating reliable modulation of individual motor units. Although isolating motoneurons was more challenging for the motor-complete group (AIS A/B; Fig. 4E), their ability to control units independently improved significantly across sessions (Fig. 5E). These findings suggest that even after severe spinal cord injury, individuals can engage residual supraspinal pathways to control spinal motoneurons (*6*, *21*) and that targeted training can enhance this control.

### Usability and Applications

These findings establish that robust motor unit–based HCI control is achievable across a wide range of impairment levels, with implementation enabled by embedded calibration and HID compatibility. Participants in this study had moderate to severe impairment in their hand function (most had finger MMT scores <=1; see Fig. S1), yet they were able to effectively don the system independently or with minimal support. Survey data indicated strong perceived usability and comfort, highlighting the potential for deploying this device for home use. Notably, motor unit activity was comparable between independently donned and assisted sessions, underscoring the robustness of the system’s motor unit detection pipeline. While the protocol was not designed to promote skill learning, participants showed clear improvements across sessions in both discrete (CHR) and continuous control (1D and 2D) paradigms, demonstrating the feasibility of sEMG-based control with a portable sEMG wristband.

Beyond the main study tasks, participants also tried using motor unit control to play video games. We adapted open-source games to work with the wristband’s discrete and continuous control signals. For example, in *Blastly*, players moved a character up and down by adjusting motor unit firing rates. In *MUShooter*, two motor units were mapped to left and right button presses for quick directional changes. *SuperTuxKart*, a racing game, required up to three motor units to steer and accelerate, and the participant used two wristbands, one on each arm, to play this game. The system also supported other controllers, such as joysticks or multiple wristbands for multiplayer gaming. Finally, we demonstrated integration with virtual reality, combining motor unit control with eye tracking for everyday tasks like dishwashing. These examples show that sEMG wristbands can fit seamlessly into daily activities and entertainment, offering new ways to improve independence and quality of life for people with tetraplegia.

### Limitations and Future Directions

The current study focused on a relatively small cohort of participants with cervical-level SCI, and while clear improvements were observed across sessions, the protocol was not optimized for skill learning. Future work should investigate the long-term learning potential and adaptation mechanisms in larger, more diverse populations to better understand the scalability of motor unit control training for practical applications. The current system requires careful calibration and placement, though the demonstrated robustness and comparable performance in independently donned sessions suggest this can be streamlined for home deployment. Key areas for development include automated calibration procedures, improved device donning mechanisms, and adaptive algorithms that can learn from user behavior to reduce setup complexity.

While this work focuses on explicit motor unit control, such decomposition may not be strictly necessary for robust interface operation. Future approaches could leverage end-to-end models that operate directly on raw or processed sEMG signals, learning motor unit-like representations implicitly to achieve stable and high-performance control. Conducting analyses that quantify the relationship between identified motor unit activity and global sEMG features could help bridge these paradigms and guide the development of hybrid systems that balance physiological interpretability with computational efficiency. Overall, the framework established here provides a foundation for optimizing motor unit selection and interface design based on individual capabilities, supporting the development of consumer-grade technologies that can be deployed with minimal technical oversight.

### Conclusion

This study represents the first demonstration of performant and volitional motor unit control in individuals with SCI using a fully wearable sEMG device. Our results confirm and extend prior observations that volitional motor unit control persists below the lesion level even in motor complete SCI, while establishing steady-state firing rate (SSFR) as a key predictor of control in tasks requiring the continuous modulation of motor unit firing rates. The AIS grade differences observed in 2D control correlate directly with the cumulative integrated firing time (CIFT) of the selected motor unit pairs. This suggests that reduced individual motor unit control independence in higher AIS grades likely arises from the limited number of motor units available below the injury level and, by extension, the associated reduction of independent sources of drive to those motor units. These constraints help explain why control performance varies by injury severity and provide both physiological insight and practical guidance for designing effective interfaces. These findings support the development of robust, user-friendly motor unit-based interfaces that can be deployed across the full spectrum of spinal cord injury severity.

## MATERIALS AND METHODS

### Surface electromyography wristband

Surface electromyography (sEMG) signals were recorded using a dry-electrode, multichannel research device, developed by Reality Labs at Meta (*28*). The device consists of two main components: a digital compute module and an analog wristband, which together enable high-fidelity sEMG data acquisition while allowing for easy donning and doffing at the wrist.

The wristband is equipped with 48 gold-plated dry electrodes, arranged into 16 bipolar channels with the sensing axis aligned along the forearm’s proximal–distal direction. The sEMG signals were sampled at 2000 Hz, bandpass filtered (20-850 Hz), digitized, and streamed via Bluetooth to a receiver dongle connected to a computer via USB. To accommodate different user morphologies, devices were available in four sizes (Table 1). Consistency in the orientation of the wristband was maintained by aligning the cinch (Fig. 1A, *Gap Location*) with the ulnar bone by palpating along the arm while sliding the wristband to the preferred location along the projection from the lateral epicondyle to the ulnar styloid process. A tape measure was used to ensure consistent placement across sessions, matching the optimal location found during the screening session.

### Screening session

An initial screening was performed using to test for detectable myoelectric activity in the forearm using a high-density surface electromyography (HD-sEMG). A group of four, 64-channel HD-sEMG grids (4mm diameter, 8.75mm pitch; Artinis TMSiSAGA64) was placed on the forearm on the participant’s self-selected (often dominant) side. During the first phase of this screening, participants were prompted for 10 sets of 5-second alternating REST and GO repetitions for all gestures from a fixed set comprising flexion, extension, radial and ulnar deviation of the wrist; supination/pronation of the arm; and flexion and extension of each digit individually. During the initial phase, experimenters visually observed modulated activity from the array electrodes during the task, noting which grids were often modulated during flexion or extension-related gestures. In the second phase, a subset of gestures (typically wrist extension and/or wrist flexion) were used during a ramp-and-hold task to 10%, 30% or 50% of maximum voluntary effort, defined using the mean power envelope from extensor or flexor channels on the grids noted by experimenters during the session. This aggregate RMS-driven approach provided effort-level visual feedback to the subject about their extensor or flexor motor drive by scaling the height of the cursor on the left edge of the screen. The intended effort ramp was provided as instruction by scrolling a visible boundary region from the right side of the screen to the left incrementally with each frame refresh, such that the stationary cursor appeared to travel “along” the path of the boundary. Motor units were decomposed offline following the session, and the location and gesture-association of the decomposed units were used to inform the gesture subset used when prompting subjects in the subsequent band placement session. All participants passed this screening step.

Following the initial screening phase, we identified the optimal wristband placement for each participant. This placement was defined as the forearm location that yielded the greatest number of active motor units and the largest set of reliably detectable gestures, and it was used consistently across all subsequent testing sessions (see Fig. E2). To localize the optimal placement, the wristband was positioned sequentially at multiple locations along the forearm, spanning from the ulnar head at the wrist to the lateral epicondyle at the elbow. At each location, wrist circumference was measured to determine appropriate wristband sizing before placement (see Fig. E2A and Table 1). Once the wristband was positioned, signal quality was first verified, after which participants were prompted to perform a set of gestures. This procedure was repeated at each location, progressing from proximal (elbow) to distal (wrist) sites. During postprocessing, placement selection prioritized the number of robustly detectable gestures and the yield of well-isolated motor units per gesture. Although distal placements (e.g., P4) sometimes produced higher motor unit counts (Fig. E2B), these increases were typically associated with greater gross movement and muscle recruitment, which participants reported as more physically demanding and less sustainable over time. As shown in Fig. E2B-C, optimal placement did not always coincide with maximal motor unit yield alone. Participant feedback regarding effort, comfort, and correspondence between neural activity and intended gestures was therefore incorporated. For longitudinal testing, placements favored locations supporting stable, low-fatigue gesture execution with high-quality, go-prompt–modulated motor unit activity; consequently, MCP01, MCP04, MCP05, MCP06, and MCP08 used P2, while MCP07 and MCP09 used P1.

### Calibration

A calibration procedure was performed at the start of each session and consisted of three stages: *Signal inspection, Motor unit discovery, and Spike control assessment* (see Supplementary Movie 1). During *Signal inspection*, participants attempted prompted gestures while the experimenter monitored live sEMG to verify task-related modulation. *Motor unit discovery* trials then identified a single motor unit template per round, selected based on waveform quality and reliable activation during “GO” epochs with suppression during “REST” epochs. Finally, participants attempted to selectively activate the identified motor units to assess controllability and consistency.

#### Signal inspection

Time-amplitude plots for the 16 EMG channels were displayed on a large central monitor visible by both the subject and experimenters. Because model-fitting required a comparison of motor unit firing rates between the GO and REST epochs for each gesture prompt, participants needed to be positioned comfortably in such a way that it was possible to suppress tonic motor unit firings. Even without decomposition, these tonic firings were clearly visible in the high-pass-filtered time-amplitude EMG signals, and because the location of each time-amplitude stream was known and mapped directly to a location on the extensor or flexor side of the arm, the experimenters could provide feedback to subjects about patterns of tonic observed activity. For example, tonic firing of a single motor unit could often be alleviated by slightly repositioning the angle at the wrist by removing or providing an additional rolled blanket to slightly change the resting flexion at the wrist. This adjustment period was restricted to no longer than 15 minutes to ensure the remaining procedures could be completed within the 4-hour time block allocated for experiments.

#### Motor unit discovery

The motor unit discovery process includes three steps: *Channel isolation, Model fitting, and Motor unit identification*, which are performed while participants are prompted to alternate between rest and attempted movement. Detailed descriptions of these steps are provided in the Supplementary Materials and Methods section, and Supplementary Movie 1 illustrates the process through representative examples.

Participants were verbally instructed to perform a specific gesture (e.g., index flexion, wrist pronation) during the prompted movement cues. Upon confirming readiness, participants followed alternating 5-second rest and 10-second movement epochs guided by audiovisual cues generated by the training software.

During channel isolation, unitary spike events were detected across all 16 sEMG channels by applying a fixed threshold of 6 standard deviations to the whitened waveforms. Each detected motor unit action potential (MUAP) was characterized by its spatiotemporal waveform. Candidate MUAPs were visualized as points embedded on a unit disk representing a cross-section of the forearm, with wristband electrodes arranged around the perimeter. The angular position of each MUAP embedding was computed as a weighted average of electrode locations, weighted by the L2 norm of the MUAP across channels. The embedding radius (ρ) was derived from the MUAP L2 norm using a logistic transform, emphasizing channel locations yielding higher-amplitude and more isolated waveforms.

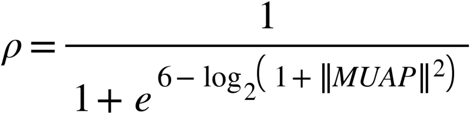

Each MUAP was briefly displayed using the polar coordinate pairs (ρ*_i_*, θ*_i_*) as a point on the disk for 1 s before disappearing. The electrode nearest the angle associated with each waveform was used to assign the selected channel for that waveform. Channel selection was not based on a fixed number of waveforms per unit time, but instead required sustained spiking activity over multiple seconds, enforced through a normalized accumulation criterion that effectively imposed a minimum firing rate (∼5 Hz) and duration (∼10 s) of consistent activity. The source location for each unsorted spike was projected as a point on a unit disk representing the cross-section of the forearm, with the wristband electrodes around the perimeter. Participants viewed a live display of the unsorted spiking activity and were encouraged to confine the activity to a localized region of the disk, corresponding to a single electrode channel. Sixteen peripheral indicators representing the electrode locations (*E_C_*), and each contained a color-mapped indicator (*F_C_*) that provided real-time feedback of channel-specific spiking activity. A channel was considered isolated once activity became sufficiently localized, at which point that channel and its immediate neighbors were selected for motor unit decomposition.

During the *Model fitting phase* for motor unit decomposition, participants continued alternating between rest and movement epochs. Visual feedback was restricted to spikes detected on the selected channel, and auditory feedback was contingent on task compliance. Spiking during rest epochs triggered corrective visual and auditory cues, whereas appropriate activity during movement epochs produced positive auditory feedback. During movement epochs, firing rates were required to remain within an acceptable range defined relative to the channel’s estimated maximum firing rate; sustained spiking that exceeded this range, indicative of excessive effort rather than selective control, triggered visual warnings and feedback encouraging reduced effort.

After three rest–movement alternations, the decomposition algorithm attempted to extract at least one motor unit meeting predefined quality and controllability criteria. If no motor units met these criteria, the oldest epochs were discarded, and the model fitting procedure was repeated, with up to three retries permitted before reverting to the channel isolation stage. If participants failed to follow prompts, which were defined as excessive spiking during rest epochs or insufficient spiking during movement epochs for more than 7 s, the procedure automatically reset to channel isolation.

Finally, during *Motor unit identification*, sEMG signals were decomposed using the MVDR-based algorithm described in the Supplementary Materials and Methods. Each successful discovery round yielded a single selected motor unit template, which was associated with the prompted gesture based on waveform quality and the unit’s ability to reliably modulate firing during “GO” epochs while remaining suppressed during “REST” epochs. Selected motor units were retained for subsequent task control.

#### Assessment

Participants were provided with feedback about the selected motor unit immediately following successful model fitting. During assessment, participants were verbally cued to alternate between REST and GO for the associated gesture, while model-classified activity was visualized as color-coded horizontal bars that scrolled from right-to-left to denote the passage of time. Transitions from REST to GO were initiated by any detected firing of the selected motor unit and were indicated by the appearance of a colored bar at the right edge of the display Conversely, transitions from GO to REST occurred when the selected motor unit remained silent for more than 200 ms, which was indicated visually by the closure of the filled rectangle at the right edge of the screen. Activity was displayed within a fixed-duration temporal window, such that sustained firing during GO caused the rectangle to progressively fill the horizontal extent of the display, while prolonged inactivity resulted in no gesture-keyed rectangle being shown. After fitting the first two gestures, participants were verbally prompted to alternate between the corresponding rectangle colors. In most cases, participants were able to reliably switch between gestures, demonstrating selective and repeatable control of the associated motor units.

### Online Firing Rate Estimation

Motor unit firing rates were computed in real-time using an online rate estimation algorithm that converted discrete spike events into adaptively smoothed continuous rate estimates (*S*). The estimator was updated at 20 Hz (Δt = 50 ms). At each update, the number of spikes that occurred in the preceding 1s window was counted and divided by the window duration to yield a boxcar firing rate. This raw rate (*R*) was then smoothed using an adaptive One-Euro filter (*29*). The filter applied a first-order low-pass with a time constant (τ) that decreased when the estimated rate changed rapidly, allowing fast responses to sudden changes while suppressing noise during steady periods. The estimator parameters and methods for updating them are described in the Adaptive Filtering section of the Supplementary Materials and Methods section.

During online cursor control tasks, the adaptively smoothed rates were received by the task software at 50ms intervals, but the core task running in Unity generated updates at 16.7 ms intervals (60 Hz refresh rate). To account for this temporal mismatch, the most recent rate estimate (*S*) was used to update the cursor’s destination. The cursor position was then interpolated frame-by-frame as a weighted fraction between its current and target location to ensure smooth, visually continuous motion of the cursor despite the asynchronous timing between the neural input updates and the Unity rendering loop.

### Motor unit control tasks

#### Click-Hold-Release (CHR) task

Participants completed a computer-based task designed to measure response latencies and motor control consistency during discrete click-and-release actions. A *click* action was asserted by activating the assigned motor unit, and deactivating the motor unit triggered the *release* action. The task consisted of repeated trials structured as a finite-state machine, updated at the display’s frame rate (60 Hz), with transitions governed by internal timers and user input sampled via keyboard, touchscreen, or gamepad. Outputs from the wristband were remapped such that they were registered in the task using the same input-handling as inputs from a standard universal-serial bus-compliant keyboard. During the mapping procedure, a sensitivity slider was set to the maximum sensitivity such that any motor unit model associated with a mapped channel caused a click event lasting a minimum of 200 milliseconds for the corresponding mapped button. If an inter-spike interval for a mapped channel exceeded 200 milliseconds, a transient release event was registered.

Each trial of the CHR task began with a REST state, during which participants were instructed to maintain a relaxed posture without clicking; the duration of the REST interval was varied randomly. Following a successful REST interval, a *click* cue appeared as a blue square on the left-center portion of the screen, signaling participants to activate the motor unit assigned to the task. Upon detection of the input click event, the time was recorded, and the color of the square changed to green as confirmation of the registered change of input state. After a short, randomized hold period (HOLD state), a second, red square was presented as the *Release cue* in the right-center portion of the screen (RELEASE state), prompting participants to deactivate the motor unit and return to the REST state. The release time was recorded. Early clicks (i.e., generated before the *Click cue*), or early releases generated before the *Release cue,* triggered an error state, indicated visually with yellow squares in place of both the *click* and *release cue* locations.

Participants were instructed to attempt to follow the *Click* and *Release* cues as quickly as possible without over-exertion. Following each session, we shared a “leaderboard” with participants, which indicated the mean latencies from the completed session as well as the best latencies across participants and an option to display past performances specific to the participant. All participants completed at least 30 trials of the CHR task using the EMG wristband. For comparison, participants also completed at least one session using a touchscreen tablet. The CHR Task States section of the Supplementary Materials and Methods and Supplementary Movie 2 demonstrates the CHR task with examples.

#### 1D Cursor Positioning Task

Participants performed a one-dimensional (1D) motor unit control task, developed in Unity (Unity Editor 2022, Unity Technologies, Inc.). In the task, participants modulated the firing rate of a selected motor unit to control the position of a cursor to reach targets positioned along a horizontal axis. The target size and distance of the target from the baseline were varied to create four target conditions. Two target distances (near and far) and sizes (small and large) were used.

At the start of each session, a brief calibration procedure was performed to determine the parameters of the function that mapped motor unit firing rate to cursor position, measured in Unity Units (UU) and referenced to a starting location at the center of the screen. The position mapping function is:

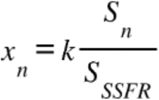

where *x_n_* is the position of the cursor at time *n* based on the smoothed firing rate (*S_n_*), which is normalized by the steady-state firing rate (*S_SSFR_*). The *S_SSFR_* is the maximum firing rate that the participant can sustain for at least one second and was measured as participants attempted to generate and hold a maximum voluntary contraction while viewing a live display of the motor unit firing rate. The gain factor (*k*) scaled the normalized firing rate to task coordinates (*UU*). The default value for *k* was 17UU, but the gain factor was occasionally adjusted manually at the start of a run to prevent participants from overexerting in reaching the far targets.

Each trial began with the cursor at a resting position at the center of the screen. Participants were instructed to volitionally modulate firing rates to move the cursor to the target region and maintain it there for a fixed duration (1s) to complete the trial. Trials were classified as *successful* if the target was reached and held for the required duration. Trials were classified as *unsuccessful* if the participant failed to acquire the target within 15 s. Audible cues indicating success or failure accompanied each attempt. Performance metrics included success rate (percentage of completed trials) and time to completion (trial duration). Supplementary Movie 3 illustrates the task with examples.

#### 2D Cursor Positioning Task

For gesture combinations that achieved a success rate of at least 80% success rate in the 1D task within a session, participants were additionally prompted to perform a similarly structured 2D task variant. This variant extended control to both horizontal and vertical screen axes. In cases where the full set of qualifying gesture combinations could not be tested within the allotted session time, experimenters prioritized combinations of greatest interest, including those suggested by participants based on gestures they felt offered their best discriminability and performance.

Each trial began with the pointer positioned at the bottom-left corner of the screen. Control inputs were normalized and scaled using the mapping used in the 1D task variant from the same session, unless otherwise specified. Gesture-to-axis assignments (horizontal vs. vertical) were determined based on verbal confirmation of participant preference.

### Task Performance Metrics

#### Path Efficiency

We quantified how closely participants followed a straight-line optimal movement of the cursor toward the target during the 2D task as the path efficiency. For each trial, we computed the path length (*PL*) as the sum of 2D Euclidean distances between consecutive cursor samples along the truncated trajectory, ending at the point of initial target collision (first contact). The straight-line distance (*SLD*) was defined as the Euclidean distance from the movement start position to the target center projected along the target axis. Path efficiency was calculated as:

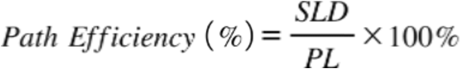

To examine within-session changes in movement strategy, trials were ordered chronologically and divided into quartiles. The first 25% of trials (Q1) represented early session performance, and the last 25% of trials (Q4) represented late-session performance. Mean path efficiency was computed separately for Q1 and Q4 for each participant and was used to quantify within-session improvement or decline.

#### Motor unit independence metric

To quantify the degree of independent control between pairs of motor units during task performance, we computed the *Cumulative Independent Firing Time* (CIFT) (*13*). For each trial, the smoothed firing-rate trajectories (*S*) for each motor unit in the pair were used to construct vectors of length *m*, corresponding to the number of samples from the time of trial onset to completion, sampled at 200 Hz. Thus, the firing rate vector for unit *j* is *S_j_* = [*S_1,j_*, …, *S_m,j_*], yielding an *m* x 2 matrix containing the firing rate trajectories for the motor unit pair.

A unit was considered active when its smoothed firing rate (*S_n,j_*) exceeded 1 spike/sec, and independently active when it was above this threshold while the other unit remained below the threshold. For each unit, the CIFT value was defined as the fraction of its total active samples during which it was independently active. Trials in which only one unit was active were assigned a value of 1. Trials in which neither unit was active were omitted. Often, both units were active during some phase of a trial; in these cases, we used the mean CIFT of each unit.

### Statistics

All analyses were performed using Python (version 3.8+) or MATLAB (R2024b) with custom analysis pipelines designed to accommodate the small sample size (N=7 participants) and non-parametric nature of the data.

We performed a nonparametric two-sample bootstrap hypothesis test for each group comparison, as follows. For each group, 10,000 bootstrap replicates of the sample mean were generated, and the distribution of the bootstrap mean-difference statistic was centered to obtain a bootstrap approximation of the null distribution. A two-tailed p-value was obtained by comparing the observed mean difference to the centered bootstrap distribution. Unless otherwise indicated, average values are presented as the mean and [2.5, 97.5] percentile of the N=10,000 bootstrap resampled estimator.

Regression modeling was performed to examine the effects of firing rate properties on performance. All fitted model coefficients are reported as the coefficient values ± standard error. Detailed descriptions of the regression methods are provided in the Supplementary Materials and Methods section.

## Supplementary Materials

## SUPPLEMENTARY MATERIALS AND METHODS

### Motor Unit Discovery - Channel Isolation

The motor unit discovery process begins with identifying the wristband channel detecting the largest sEMG signal while participants formed prompted gestures. Only the selected channel and nearest neighbors are used during motor unit decomposition to detect motor unit action potentials. During the channel isolation step, participants are prompted to perform a specific movement and restrict their motor unit activity to a localized region of the arm. Spike detection criteria required that the absolute value of the sEMG at the center location on at least one channel exceed a constant threshold (6 standard deviations), and for the peak of the summed absolute sEMG activity to occur in the temporal center of the 10-ms window. Putative MUAP waveforms were associated with the channel maximizing the peak absolute value of sEMG in the window.

To determine in aggregate which channel subset would be used during the motor unit model-fitting phase, candidate-spike counts were smoothed to produce rolling threshold-crossing rate estimates (*R_C,m_*) in 1-second windows (*m*) for each channel (*C*) using a stride of 5-ms. The maximum windowed rate from each channel was then estimated over a rolling window of 5 min. In parallel, a cumulative spike count (*N_C_*) generated on each channel during movement epochs was updated on the same timescale. Normalized channel activations (*A_C_*) were then estimated using the following equation:

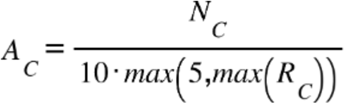

where the constant values of 10 and 5 correspond to a minimum duration of sustained spiking activity and a lower bound on the maximum allowed estimator for the per-channel firing rates, respectively.

While performing the prompted gestures, participants received visual feedback showing the approximate location of the detected motor unit activity, which was mapped to the interior of a unit disc representing the cross-section of the arm and wristband. The 16 electrode locations (*E_C_*) were displayed around the perimeter, and color-mapped concentric circular indicators (*F_C_*) were used to deliver real-time user feedback. Values of *A_C_* were rendered visually by updating the radius of *F_C_* with each refresh of the monitor frame. As values of *A_C_* approached 1, the radius of the concentric, colored circle *F_C_* filled the circle for the corresponding electrode (*E_C_*). A channel is selected when *A_C_* exceeds a value of 1, indicating sustained, movement-specific activity. Following channel selection, the discovery procedure advanced to the sEMG Decomposition model-fitting stage.

### Motor Unit Discovery - sEMG Decomposition Model Fitting

As in the *Channel Isolation* step, the sEMG decomposition models were fit while the participants alternated between 5-second rest and 10-second movement epochs for the instructed gesture. The model-fitting algorithm operates through three iterative steps: (1) motor unit identification by detecting and clustering candidate MUAPs using signals from the selected channel subset to estimate unique templates, (2) spike time inference using Minimum Variance Distortionless Response (MVDR) template-match filtering, and (3) iterative peeling to subtract identified MUAPs and reveal additional motor units.

#### Motor Unit Identification

MUAPs are characterized by a singular peak having a large amplitude relative to the background signal. Thus, local peaks in the sEMG were identified as MUAP candidates if they were local maxima within a 20-ms window and exceeded the 50th percentile of the absolute values of sEMG. A peak detector algorithm was applied to each channel independently and detected MUAP candidates, which were represented as tuples (*C*_i_, *t_i_*), indicating the channel number and time of detection.

A “MUAP snapshot” is created for each candidate MUAP by capturing a spatiotemporal window of sEMG signals that includes a 12 ms snippet of sEMG centered on *t_i_* for the selected channel (*C_i_*) and its nearest neighbors (*C_i-1,Ci+1_*). Thus, the MUAP snapshot is a 3x24 matrix of sEMG samples.

K-means clustering (*K* = 10) is applied to each group of MUAP candidates, which are the resultant 3x24xn candidate waveforms recovered from the cumulative *Channel Isolation* rest/go iterations, once a channel has been selected. MUAP templates are then calculated by averaging the MUAP snapshots within each cluster. Clusters are disregarded if the signal-to-noise ratio (SNR) of their corresponding MUAP template is low. The SNR of a MUAP template is defined as the difference between its minimum and maximum values, divided by the sEMG noise level, which is represented by the median-absolute-deviation (MAD)-based standard deviation of sEMG. MUAP candidates are removed from a cluster if their peak location differs from the peak location of the MUAP template. Clusters with fewer than 10 assigned MUAPs are removed, followed by an iterative merging procedure in which the two clusters with the most similar MUAP templates are merged. Similarity is determined by the Euclidean distance between two MUAP templates, normalized by the average L2 norm of the MUAP templates. This merging process continues until no two pairs have a normalized distance less than 0.05.

Following this k-means procedure, clusters are expected to represent motor units. However, because not all clusters are clearly isolated, only those with a large mean, a small standard deviation, and many individual members are selected for the next steps. A metric (*M*) that combines these statistics, expressed as follows, is used to select motor units:

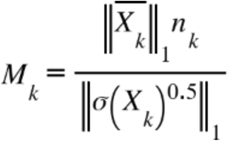

where:

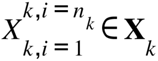 is the set of *n_k_* snapshots of MUAPs assigned to the cluster *k* and has *T* rows and *C* columns, where *T* is the temporal window (*T* = 24 samples), and *C* is the number of channels (*C* = 3).

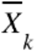 is the median and σ(*X_k_*) is the variance of MUAP snapshots, both having the same size as 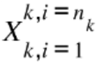.

#### Spike Time Inference

Minimum Variance Distortionless Response (MVDR) filters were used to detect the trigger times of given templates from noise. MVDR filters have a closed-loop solution and are computed as follows:

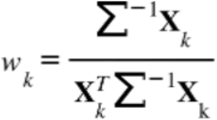

where *w_k_* is the MVDR filter of the motor unit *k*. Σ is the noise variance in sEMG, and **X***_k_* is the MUAP template of the motor unit *k*. Due to its normalization factor, the MVDR filtered data should exhibit a peak with a magnitude close to 1 when a MUAP is present in the data. Therefore, after applying the MVDR filter to sEMG, peak times greater than 0.7 are considered spike times. If multiple motor units are selected, they are ordered from largest to smallest based on their L2 norm and applied sequentially. After inferring the spike times of one motor unit, the motor unit is peeled off as described next. The MVDR filter of the next motor unit is then applied to the residual sEMG using the same method.

#### Iterative Peeling

After inferring the spike times of a selected motor unit, the motor unit is “peeled off” from the sEMG signal. This process involves fitting a scaled version of the MUAP template at each inferred spike time and then subtracting it from the sEMG. The scaling factor for each MUAP instance is estimated independently to account for amplitude variability. This factor is chosen to minimize the difference between the observed MUAP instance and the MUAP template. However, if the estimated scaling factor falls outside the range of 0.9 to 1.1, it suggests potential corruption of the MUAP shape. In such cases, the MUAP template is subtracted without scaling.

The precision of inferred spike times, limited by the sEMG sampling rate, can introduce artifacts in the residual due to even minor discrepancies. To mitigate these artifacts, the sEMG is upsampled by 10-fold within local temporal windows surrounding the inferred spike times and spike times are refined to a finer temporal resolution. After subtracting all inferred MUAPs from the sEMG, the remaining residual undergoes the same process to identify subsequent sets of motor units and their spike times. This iterative process continues until a maximum of 10 motor units are identified, or until no further motor units can be found.

#### Motor Unit Model-Selection

In addition to the signal quality and template-match metrics, motor units were filtered based on a controllability metric, *C*, which required that any selected unit maintain an average firing rate ≥2 spikes/s during movement epochs. The controllability metric is defined as

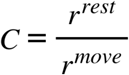

where,

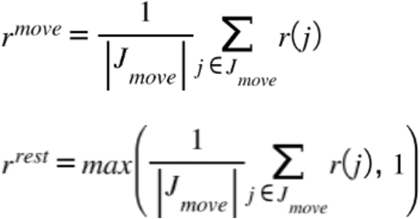

for the non-smoothed rate estimates *r* obtained from counting spikes in a sliding one-second window during discrete sample update *j*. Units were only selected if *C* ≥ 2. In cases where the iterative procedure resulted in multiple motor units meeting the selection criteria, the unit with the largest value of *C* was selected to be associated with the gesture movement.

Motor unit detection models were fitted following this procedure for up to 10 motor units per session. Because the specific subset of gestures was different for each subject, we typically fitted a model for between 3 and 6 gestures per session.

### Adaptive filtering

The adaptive filter applied a first-order low-pass filter with a time constant (τ) that decreased when the estimated rate changed rapidly, allowing fast responses to sudden changes while suppressing noise during steady periods. The key estimator parameters and methods for updating them are described below:

- β = 0.05, which controls the sensitivity of the smoothing time-constant to recent fluctuations in the smoothed output
- dτ = 0.01, which sets the fixed smoothing level for smoothing the changes in the rate estimate; and
- τ_max_ = 1.0, which effectively controls the lower bound on the cutoff frequency for the adaptive smoothing lowpass filter.

The filter’s recursive update at each 50 ms step *n* was:

1. Compute the time derivative of the firing rate:

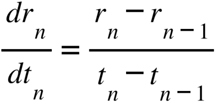

2. Lowpass filter the derivative of the adaptive rates (*ds*):

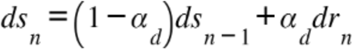

Where *α_d_* = 0.83 and is a fixed exponential smoothing coefficient.

3. Compute the adaptive time constant (τ*_n_*):

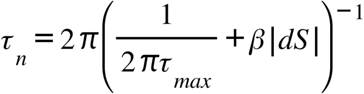

From which the minimum exponential smoothing filter cutoff is 1/(2πτ_max_) during quiescent periods, where *dS* = 0.

4. Update the smoothed rate *S_n_*:

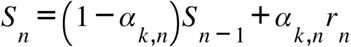

Where α*_k,n_* is the adaptive exponential smoothing coefficient for the online rate estimator *S_n_* and is calculated by:

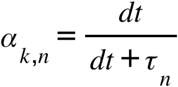

### CHR Task States

Each Click–Hold–Release (CHR) trial progressed through a fixed sequence of states, each with well-defined temporal requirements. All times were measured using high-resolution monotonic clocks (JavaScript performance.now() function). Prior to starting the task, the experimenter defined a total number of required trials to be completed for the session to register as completed in the task database. Using this target number of trials, a set of wait times drawn from a clipped exponential distribution bounded between 1000–4000 ms was sampled once, then shuffled randomly to associate the same distribution of wait and hold times to the REST and HOLD states as described below. Upon completion of a session, the results were logged on a database server, which automatically summarized the session performance. The participant was shown their performance by redirecting the browser to a “results” route following completion of the final trial, which included a “leader board” that allowed comparison of reaction times against other participants or the experimenters.

#### States -3, –2, and –1 (intertrial relaxation)

To prevent transient error state indications, which could otherwise rapidly flicker up to the frame rate of the task monitor (60 Hz), a ≥350 ms debounce state was used between trials (State -3), which always transitioned to a relaxation state (State -2). In the event of an error, a flag set before State -3 caused the appearance of yellow squares in the locations of both the *click cue* and the *release cue*. Because the participants were not always aware that they were sustaining a clicked input state, these error cues remained on the screen until the release of the click, at which point the state timer for State -2 was allowed to advance. During State-2, early click errors were possible, but required a sustained click and hold lasting at least ≥250 ms, which helped prevent confusion due to non-debounced transient clicks following a sustained, erroneous click. If no error occurred in State -2 for ≥350 ms, the task advanced to State -1, at which point there was no sustained click requirement for an early click error. If no error occurred in State -1 for ≥350 ms, the task advanced to State 0. During State -2 or State -1, an early click reset the system to State –3. A global timer, which prevented trials from exceeding 20s total, was not reset during these errors. Early click errors, which reset the trial to State -3, did not count as a completed trial, whereas a timeout on the global timer did count as a completed trial, incrementing the nominal number of trials performed against the number of trials set (default: 30 trials).

#### State 0 (REST)

Following the intertrial relaxation states, a variable REST period was drawn from the array of possible REST durations generated as described above. Participants were required to remain fully relaxed for the full duration. An early click before the cue onset immediately aborted the foreperiod and returned the task to State -3, indicating the error.

#### State 1 (CLICK)

Following REST, the *click cue* appeared, and the task entered State 1. Participants were required to click during this phase. Click latency was measured as the interval between cue onset and the rising edge of the detected click (Fig. 2A). A successful click during this phase transitioned the task to State 2. While it was technically possible for subjects to timeout during this phase, timeouts rarely occurred except for rare technical difficulties (such as a dead battery on the wireless device at the end of a session, or a parameter misconfiguration by the experimenter).

#### State 2 (HOLD)

Following CLICK, participants were instructed to hold the clicked state for a second variable delay period drawn from the array of possible HOLD durations generated as described above. Failure to sustain the click for the duration of the HOLD state was treated as an error, resetting the task to State -3. After the hold duration elapsed, the release cue appeared, and the trial transitioned to State 3.

#### State 3 (RELEASE)

At the end of the HOLD state, participants were cued to release following the presentation of a red square at the right-center of the screen. Release latency was measured from cue onset to the first detected falling edge of the input (Fig. 2A). Following a successful release, the trial counter was incremented, and the state was reset to State -3, without the flag to show the error indicators.

### Regression Modeling

#### Logistic Regression Model: 1D-Cursor Per-Trial Success by Firing Rate

To quantify how task difficulty and firing-rate normalization influenced per-trial task success (Fig. 5B), we fit a generalized linear mixed-effects model (GLME) with a binomial likelihood and logit link. The dependent variable was the trial outcome (*Succ*), encoded as 1 for successful trials and 0 otherwise. Fixed effects included an intercept term, the normalized target distance (*NormDistance*), the steady-state firing rate (*SSFR*) determined during calibration for each unit prior to the 1D cursor task, and a fixed effect interaction between *NormDistance* and *SSFR*. *NormDistance* represents the position of the target expressed in units of SSFR, correcting for session-specific gain values used to map firing rate onto screen units in the Unity environment. Since gains varied across session–gesture blocks, identical “Near” and “Far” targets on the screen corresponded to different fractions of each unit’s SSFR, making *NormDistance* the appropriate normalized predictor. The dynamic range of the motor unit firing rate, in raw spikes per second, could be a significant physiological correlate of its behavior in this task; the interaction term was therefore included to capture this influence. The full model specification was:

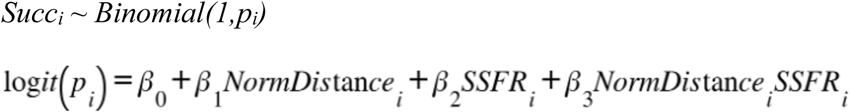

The model was fit in MATLAB (R2024b) using the fitglme function using a logit link function and binomial distribution.

This formulation allowed us to estimate population-level effects of normalized target distance on success, while accounting for heterogeneity across target sizes in both baseline difficulty and the influence of firing-rate–related variables (Fig. 5B).

#### Linear Mixed Effects Model: 2D-Cursor Trials Per Minute by CIFT

To quantify how independent motor-unit control contributed to overall throughput during the 2D-cursor task, we modeled throughput (*TrialsPerMinute*) as a function of the mean CIFT (*Mean_CIFT*) computed across all trials of each run, with a random effect grouping by subject (Fig. 5C). For every subject–session and gesture-pair combination, we calculated trial throughput as the number of completed trials divided by the total time spent attempting those trials, with unsuccessful trials assigned their full timeout duration to reflect their contribution to task time. Mean CIFT for each combination was computed by averaging CIFT across all trials (successful and unsuccessful) with missing values (from trials with no firings of either motor unit within the gesture pair) omitted.

We then fit a linear mixed-effects model with throughput as the continuous response variable, CIFT as the fixed effect, and subject-specific random intercepts to account for baseline differences in performance across participants:

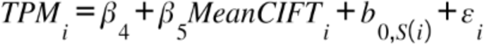

where 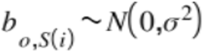 is a random intercept for subject *S*, and 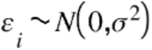 is the residual error. This model estimates the population-level increase in throughput associated with higher independent motor unit control while permitting each subject to have a distinct baseline performance level. In MATLAB (R2024b), the model was fit using the fitlme function, excluding trials for which CIFT could not be computed (i.e. when neither unit became active prior to the trial timeout).

Population-level predictions and 95% confidence intervals (fixed-effects component) were evaluated over the observed range of Mean CIFT to generate the smooth regression line and shaded confidence band shown in Fig. 5C.

#### Generalized Linear Mixed Effects Model: 2D-Cursor CIFT Longitudinal Model

To characterize how normalized target position, longitudinal training, and within-session practice influenced the proportion of within-trial time spent with only one of the two units active (Fig. 5E), we modeled the Cumulative Integrated Firing Time (CIFT) on a per-trial basis using a generalized linear mixed-effects framework. CIFT values were bounded to the open interval [0,1] and transformed using the standard normal inverse cumulative distribution function, producing a probit-scaled CIFT (*probitCIFT*) response suitable for linear mixed-effects modeling. To avoid numerical singularities in the probit transform, CIFT values of exactly 0 or 1 were mapped into the open interval [0,1] using a small constant δ = 10^-3^ using the following equation:

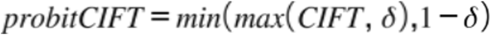

This adjustment ensures well-defined transformed values while remaining negligible relative to the empirical resolution of the CIFT measure.

The primary fixed effects included: (i) Normalized Distance (*NormDistance*), representing target position expressed relative to each unit’s steady-state firing rate; (ii) Session Index (*SessionIndex*), a continuous variable in the range [−1,1] representing each subject’s relative session number within the 2D cursor task (centered such that earlier sessions take negative values and later sessions positive values); (iii) *Severity*, a categorical factor describing neurological impairment level (classified as AIS A/B or AIS C/D); and (iv) the within-run-target index (*WithinRun*), a continuous predictor capturing the trial’s ordinal position within a gesture-pair and target–specific run, normalized to [-1, 1] (i.e. this value incremented as the sequence [1, 1, 1, 1, 2, 2, 2, 2, …] with index incrementing for each new round of targets). An interaction term between *SessionIndex* and *Severity* allowed longitudinal changes in CIFT to differ by impairment class.

To capture subject-specific heterogeneity in both baseline *CIFT* levels and learning-related progression across sessions, we included random intercepts and random slopes in the Session Index for each subject (*Subject*). This allowed the longitudinal trajectory of probit-transformed *CIFT* to vary across individuals while estimating population-level effects. Trials were excluded if *CIFT* was undefined (i.e., for trials in which neither of the selected motor unit firings maintained a non-zero value for the duration of the trial). Subsets of trials in runs where the participant was allowed to try “extra trials” (i.e., *WithinRun* > 1) were excluded from the modeled range. Similarly, subject-session combinations with “extra sessions” (i.e., *SessionIndex* > 1) were excluded from the modeled range. The final model took the form:

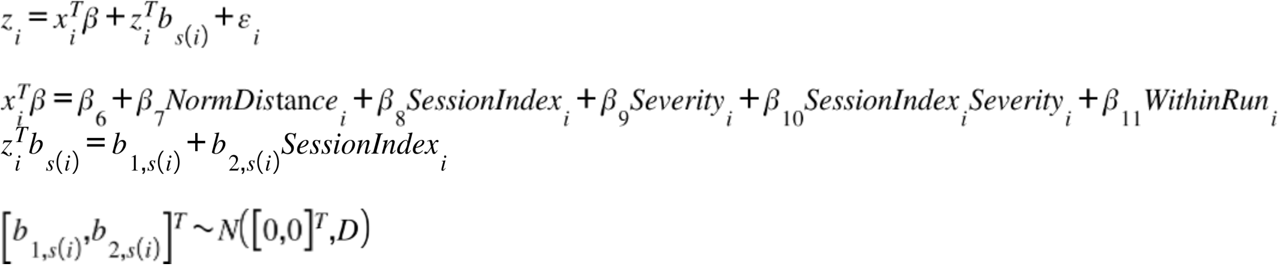

where:

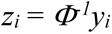 is the probit transform of the (bounded) CIFT (*y_i_*)

*x_i_* is the vector of fixed-effect predictors for trial *i*

β are the population-level coefficients

Each *z_i_* selects the random effects for the subject associated with the trial *i*

*b_s(i)_* are the random effects for subject *S*

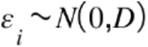 is the residual

The random effects follow 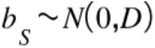 with *D* the subject-level random-effects covariance matrix. In MATLAB (R2024b), this generalized linear mixed effects model was fitted using the fitlme implementation, using the restricted maximum likelihood option for fitting the model.

This formulation enables quantification of both within-session factors (target distance, run structure) and cross-session learning effects, while accounting for differences associated with impairment level and individual variability in longitudinal trajectories (Fig. 5E).

## SUPPLEMENTARY FIGURES

**Fig. S1.**
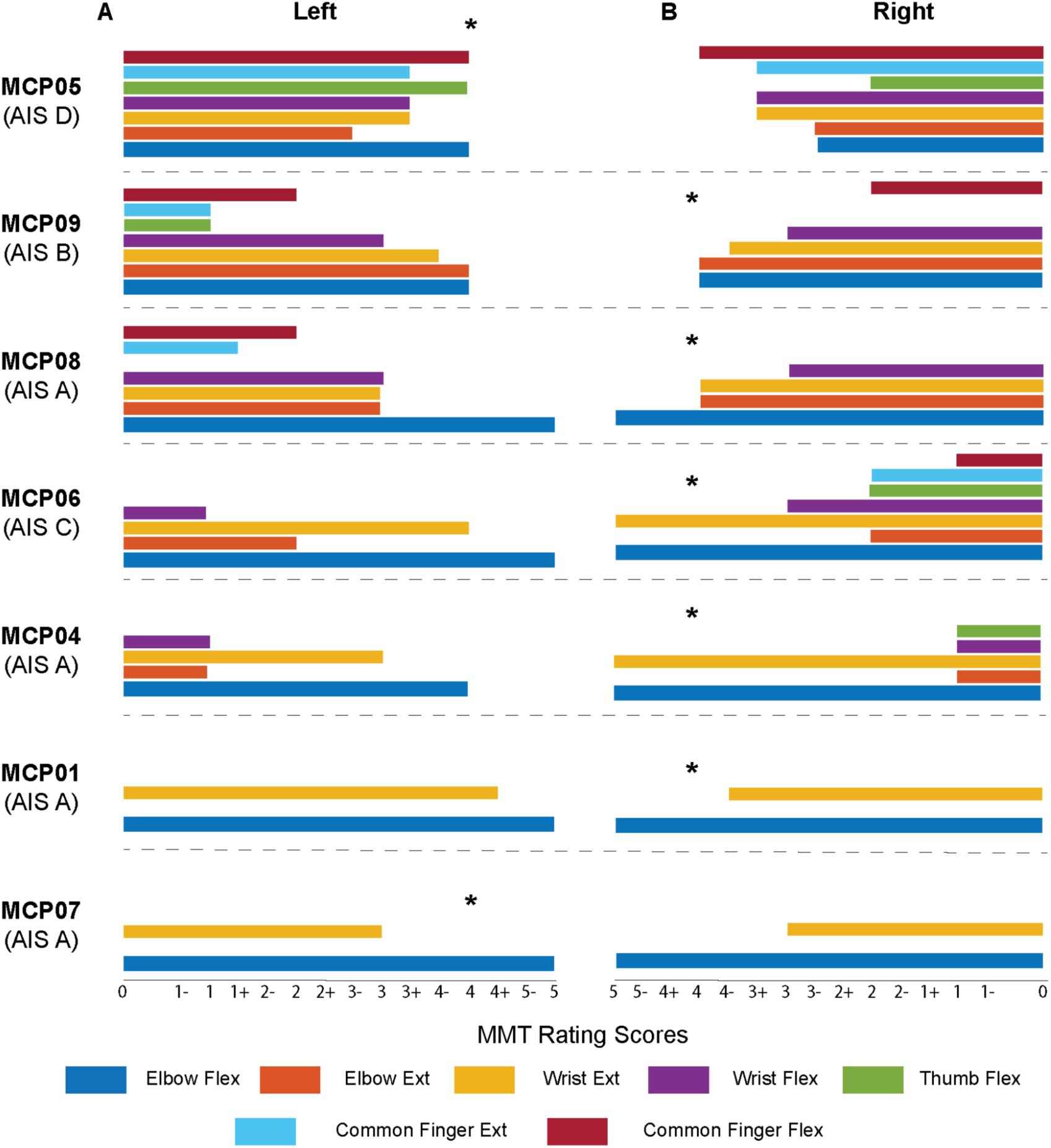
Motor impairment ratings by subject. The Manual Muscle Test (MMT) was performed by a licensed occupational therapist. Existing MMT grading systems feature a + and - in conjunction with a 0-5 score reflecting finer gradations of range of motion, with a 0 indicating no visible or palpable contraction and a 5 indicating normal strength and maximal resistance. The participants are ordered from least to most impaired, based on cumulative MMT scores, and the AIS rating for each participant is indicated in parentheses under their study ID code.

**Fig. S2.**
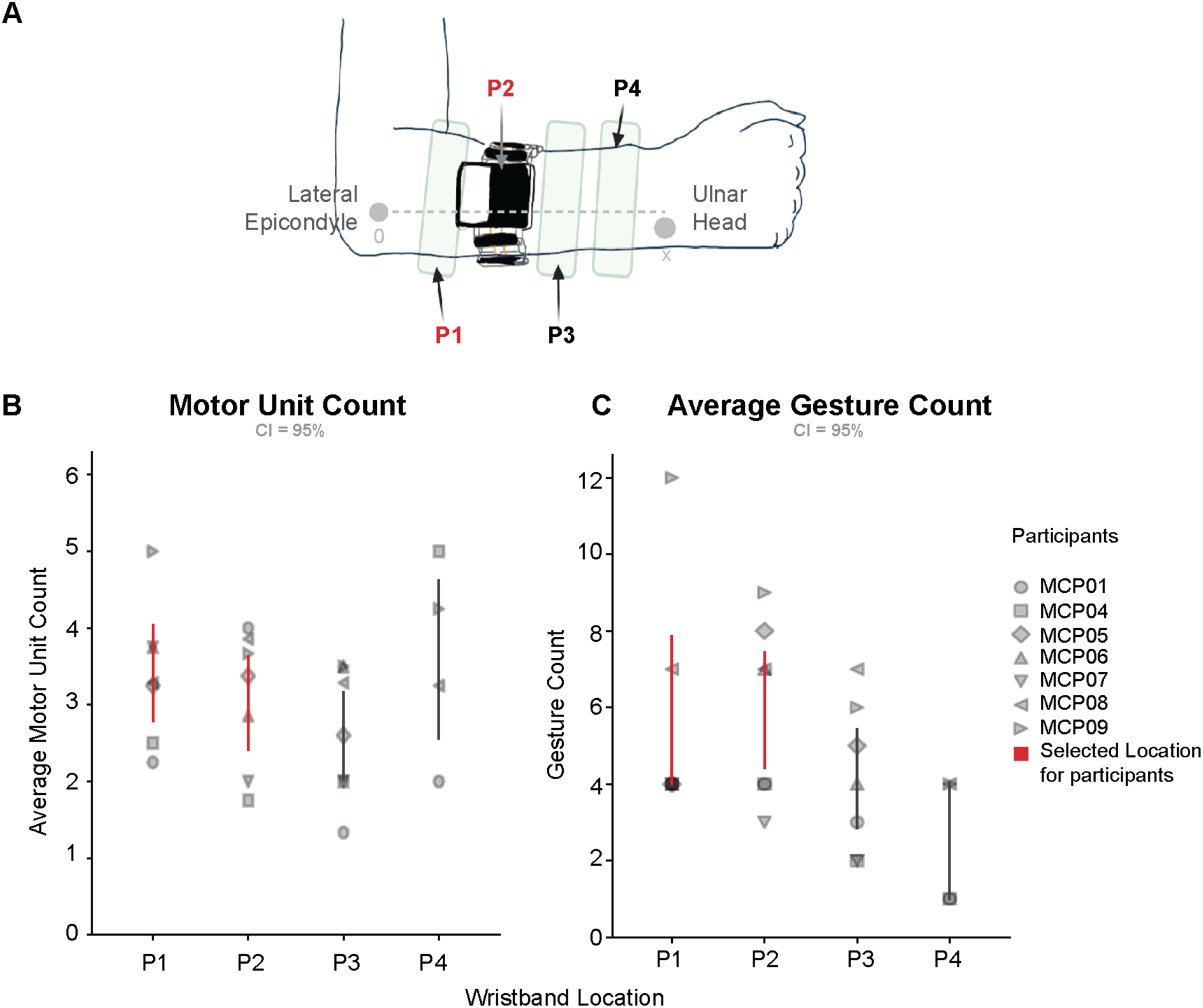
Placement of EMG wristband on the forearm. (**A**) Schematic of the forearm showing the approximate placement of the wristband and electrode positions (P1–P4) between the lateral epicondyle and the ulnar head. P1 and P2 (red labels) indicate the locations selected based on the highest motor unit counts and the greatest number of gestures detected. For longitudinal testing, MCP01, 04, 05, 06, and 08 used P2, whereas MCP07 and MCP09 used P1. (**B**) Average *motor unit count* (mean ± 95% CI) across participants and wristband locations. Each marker shape represents an individual participant; red bars denote the locations selected for optimal signal quality. Proximal sites (P1–P2) generally yielded higher motor unit activity compared to distal sites (e.g., P3, P4), which showed lower counts without limb movement. (**C**) Average *gesture count* (mean ± 95% CI) across participants and locations. A similar trend was observed, with P1–P2 supporting a greater number of reliably detected gestures. Together, these results indicate that more proximal electrode placements (P1–P2) provide superior motor unit yield and gesture detection compared to distal placements, where activation often requires gross limb movement.

**Fig. S3.**
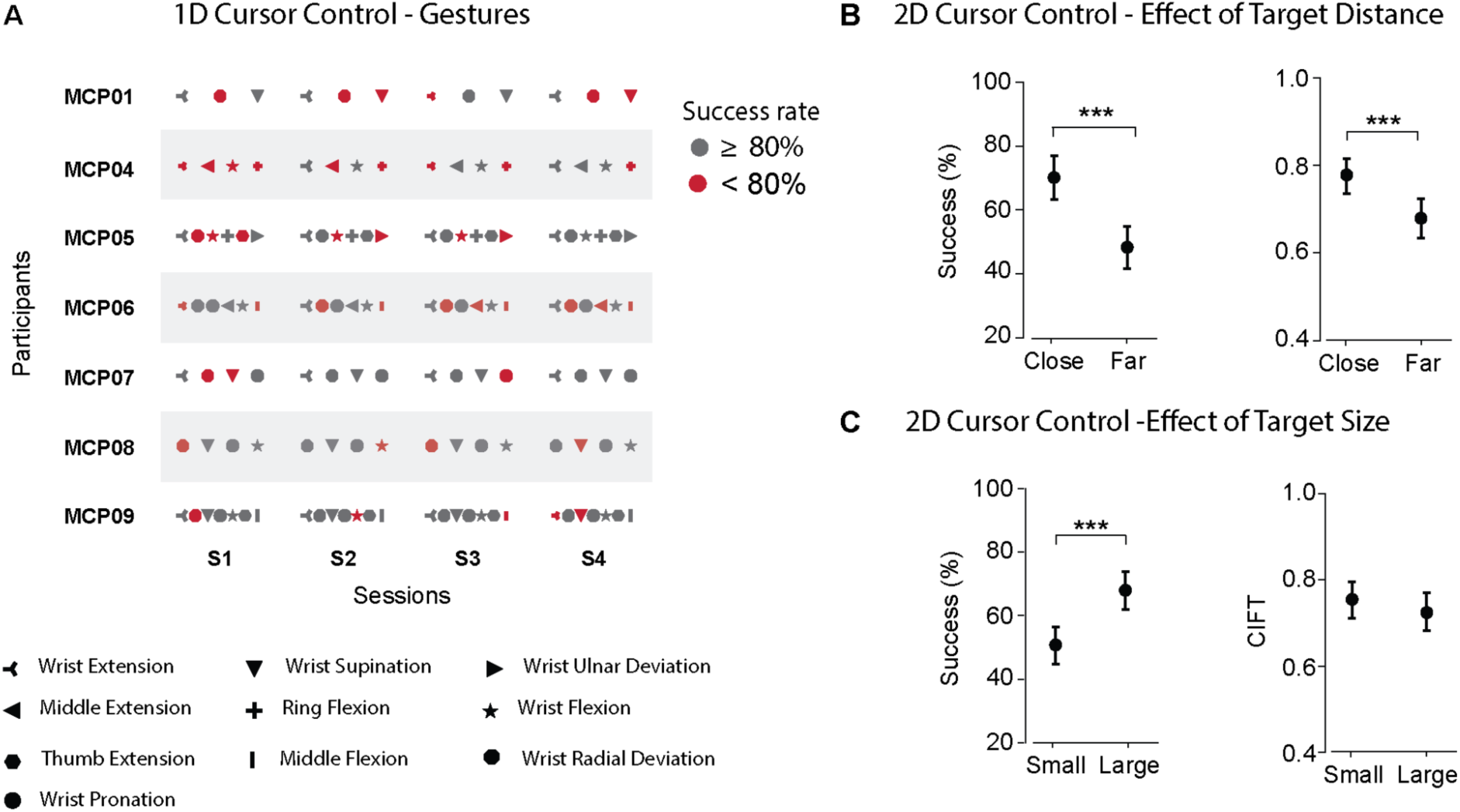
1D and 2D cursor control task. (**A**) Table presenting the gestures evaluated in the 1D cursor control task and their corresponding success rates across sessions and participants. (**B**) Quantification of mean success rate and cumulative independent firing time (CIFT) across target distance. (**C**) Quantification of mean success rate and cumulative independent firing time (CIFT) across target size. The circle marker on the plots indicates the mean value of the performance metric. All error bars indicate the 95% confidence interval of the mean performance metric computed with bootstrap N=10,000. Statistical significance was assessed with two-tail bootstrapping (N=10,000): p<0.05 (*), p<0.01 (**), p<0.001(***).

## SUPPLEMENTARY TABLES

**Table S1.** Regression Model Coefficients. Each of the statistical models, along with the corresponding subpanel from Fig. 5, has the fitted model coefficient value, standard error, and significance from all model fixed effects. Only the longitudinal model for CIFT had significant random effects, which are shown in gray with corresponding levels (subject) in parentheses. One participant (MCP07, AIS A) had a significantly larger gain in CIFT across sessions in the 2D cursor task. For the same task, a second AIS A participant (MCP08) started with a significantly lower CIFT. MCP09 (AIS B) was an outlier in that his CIFT was significantly higher at baseline; notably, this subject had a tendon transfer on the arm used in this study.

| Model | Figure | Term | Value | SE | p |
| --- | --- | --- | --- | --- | --- |
| Logistic Regression | 5b | $\beta_0$ | -0.069 | 0.452 | 0.878 |
| Logistic Regression | 5b | $\beta_1$ | -1.193 | 0.626 | 0.057 |
| Logistic Regression | 5b | $\beta_2$ | 0.076 | 0.027 | 0.005 |
| Logistic Regression | 5b | $\beta_3$ | 0.034 | 0.038 | 0.368 |
| LME Throughput vs CIFT | 5c | $\beta_4$ | 0.186 | 0.629 | 0.768 |
| LME Throughput vs CIFT | 5c | $\beta_5$ | 6.290 | 0.913 | 0.000 |
| GLME Longitudinal CIFT | 5e | $\beta_6$ | 0.613 | 0.326 | 0.061 |
| GLME Longitudinal CIFT | 5e | $\beta_7$ | 0.474 | 0.233 | 0.042 |
| GLME Longitudinal CIFT | 5e | $\beta_8$ | 0.153 | 0.091 | 0.093 |
| GLME Longitudinal CIFT | 5e | $\beta_9$ | -0.755 | 0.127 | 0.000 |
| GLME Longitudinal CIFT | 5e | $\beta_{10}$ | 1.321 | 0.598 | 0.028 |
| GLME Longitudinal CIFT | 5e | $\beta_{11}$ | -0.707 | 0.399 | 0.077 |
| GLME Longitudinal CIFT | 5e | $b_{2(s07)}$ | 0.576 | 0.278 | 0.038 |
| GLME Longitudinal CIFT | 5e | $b_{1(s08)}$ | -0.771 | 0.341 | 0.024 |
| GLME Longitudinal CIFT | 5e | $b_{1(s09)}$ | 1.015 | 0.335 | 0.002 |

## SUPPLEMENTARY MOVIES

**Movie 1: <u>Study calibration phase</u>**

**Movie 2: <u>Click Hold Release (CHR) Task</u>**

**Movie 3: <u>1D Cursor Task</u>**

**Movie 4: <u>2D Cursor Task</u>**

**Movie 5: <u>Wristband Participant Independent Use</u>**

**Movie 6: <u>Additional Developments for Wristband</u>**

## ACKNOWLEDGEMENTS

The authors thank the participants and their friends and family for their time and enthusiastic participation in the study. We are also grateful to Sam Russell for helping to create the software used in this study and to Debbie Harrington for assistance with participant recruitment and regulatory activities.

## Author contributions

Conceptualization: DDR, MM, LB, NM, EF, MB, SN, DAG, DJW

Methodology: DDR, MM, LB, NM, EF, MB

Investigation: DDR, MM, LB, KK, NV, JS, AB

Visualization: DDR, MM, LB, KK, NV, JS, MS, AG, PHS, ZA

Funding acquisition: SN, DAG, DJW

Project administration: SN, DAG, JS, JLC, DJW

Supervision: SN, DAG, SN, JLC, DAG, DJW

Writing – original draft: DDR, MM, LB, NM, EF, MB, SN, DJW

Writing – review & editing: DDR, MM, LB, NM, EF, MB, JL, JY, PW, DM, SN, KK, JS, AB, JLC, DAG, DJW

## Data and materials availability

The paper and its Supplementary Materials contain all the necessary data to assess the conclusions.

